# Sex-dependent resource defense in a nectar-feeding bat

**DOI:** 10.1101/2021.08.16.456451

**Authors:** Sabine Wintergerst, York Winter, Vladislav Nachev

**Author notes:** Institut für Biologie, Humboldt Universität, 10099 Berlin, Germany (York Winter). QUEST Center, Berlin Institute of Health at Charité – Universitätsmedizin Berlin, 10117 Germany (Vladislav Nachev).

## Abstract

Aggressive resource defense spans from the transient monopolization of a resource up to the long-term maintenance of a territory. While such interference competition is common in nectar-feeding birds, reports in nectar-feeding bats are rare. *Glossophaga* bats have been observed to temporarily defend flowers but the extent of this monopolization, its effects on nectar intake, and underlying sexual differences have remained unknown. We investigated resource defense behavior of *Glossophaga mutica* in the laboratory. We presented bats with two patches of computer-controlled artificial flowers and tracked individual nectar intake. Furthermore, we established an automated method for detecting aggressive interactions at the artificial flowers. Theoretical models of interference competition predict more aggressive interactions when resources are spatially more clumped. To test this, we varied resource distribution across two patches from clumped to dispersed and monitored bats’ interactions in one male, one female, and four mixed-sex groups. Males engaged in aggressive interactions more often than females and in each group some individuals defended clumped artificial flowers against others. Subordinate males experienced a substantial decrease in nectar intake, while females were only marginally affected by male aggression. These results suggest that aggressive interactions and their effect on nectar intake are sex-dependent in *G. mutica*. Furthermore, aggressive interactions were more frequent and resource defense was only successful when resources were clumped. Our experimental set-up allowed us to perform an automated test of models of interference competition with a mammal under controlled laboratory conditions. This approach may pave the way for similar studies with other animals.

**Lay summary:** Males bully other males to get more food, but only when food is easy to defend. When flowers are spread out nectar-feeding bats rarely engage in fights. However, when there are rich flowers in one spot and no flowers elsewhere, some males start attacking others, denying them access to the nectar. Females do not seem bothered by such male bullies, but when there are no males around, some females become bullies themselves.

## 1. Introduction

Competition for limited resources like food or mates can be indirect, by exploiting a common resource and preventing others from benefiting from it (Paton and Carpenter 1984), or direct, by aggressively defending a resource. The latter is known as interference competition (Amarasekare 2002). Aggressive resource defense by excluding competitors leads to priority of access to those resources and thus establishes dominance. One individual is dominant over another if it directs aggressive behavior towards it (chasing, threatening, biting, etc.) while receiving little or no aggression from the other (Chase et al. 2002). In the extreme, dominance behavior can lead to exclusive territoriality. Territoriality is a continuum of behaviors, from the transient monopolization of a preferred feeding opportunity to the longer-term defense of an area as exclusive territory. The rules of economic defendability (Brown 1964) determine the adaptive compromise to which a species’ dominance behavior will evolve and develop along this continuum.

The establishment of feeding territories is well known for nectar-feeding birds (Gill and Wolf 1975; Carpenter and Macmillen 1976; Boyden 1978; Ewald and Carpenter 1978). The cost of defense, however, a key parameter in the economic defendability equation, is likely much higher for a nocturnal, echolocating bat than for a diurnal, visually oriented bird. The successful resource defense is only possible after the competition is detected. Visual detection in the daylight works well over long distances. In contrast, for a nocturnal, echolocating bat, especially for phyllostomid bats that are able to echolocate with “whispering” or narrowly-focused calls (Howell 1974; Gonzalez-Terrazas et al. 2016; Yoh et al. 2020), detecting intruders at a feeding territory’s boundary would require expensive patrolling flights.

Within bats, the flower visitors have an advantage in establishing territories compared to insect-hunting bats, because bats feeding on insects must continually scan for elusive prey by active echolocation, whereas flower visitors can approach a target with minimal echolocation when seeking specific flowers at known locations (Thiele and Winter 2005; Winter and Stich 2005; Rose et al. 2016; Gonzalez-Terrazas et al. 2016; Simon et al. 2021). Thus, it is not surprising that the longer-term defense of extensive feeding territories as commonly observed in nectar-feeding birds is not known for glossophagine, nectar-feeding bats (but see Watzke 2006 for nectar-feeding flying foxes). Nonetheless, several observations have documented aggressive food defense by glossophagine bats. The inflorescences of *Agave desmettiana* with their copious nectar (Lemke 1985) may be defended by males or females of *Glossophaga soricina* against conspecifics but only during some hours of the night (Lemke 1984, 1985). When left unguarded, intruders quickly exploit the opportunity to feed from the previously defended plants. The Costa Rican bat *Glossophaga commissarisi* occasionally defends and temporarily monopolizes single inflorescences of the understory palm *Calyptrogyne ghiesbreghtiana* against other hovering bats, perching bats and katydids (Tschapka 2003). A commonality in these reports is that the defense does not cover the area of a typical feeding range but is restricted to a single or a few flowering plants and is also limited to a small number of hours during the night. While these reports demonstrate that glossophagine bats can show aggressive resource defense, the extent of territoriality and its relation to sex remain unknown.

In this study, we investigated for a nocturnal, nectar-feeding mammal, the flower-visiting bat *Glossophaga mutica* (previously *G. soricina*, Calahorra-Oliart et al. (2021)), the role of aggressive interactions for gaining access to nectar food. Economic defensibility predicts that clumped resources will be easier and less costly to defend (Brown 1964; Gill and Wolf 1975; Carpenter and Macmillen 1976). We tested this predictions using a naturalistic foraging paradigm in the laboratory. The occurrence of resource defense is predicted to be highest at intermediate levels of food abundance (Gill and Wolf 1975; Carpenter and Macmillen 1976; Carpenter 1978; Grant et al. 2002). In line with this prediction, the transient nature of nightly defense behavior observed in the field suggests that changes in food-abundance or food-requirements that occur within the night affected the strength of the observed behavior. To mimic the natural situation of chiropterophilous flowers, which often replenish their nectar more or less continuously throughout a night (e.g. Tschapka and von Helversen 2007), we programmed artificial flowers to provide nectar with a fixed interval reward schedule. Once a nectar reward had been taken by any bat, the fixed interval had to pass before the next reward was available at this flower. Theoretical models of interference competition predict that clumped resources lead to more agonistic behavior and resource defense than evenly dispersed resources (Brown 1964; Carpenter and Macmillen 1976; Grant 1993). To include a test of this prediction in our experimental design, we spatially subdivided our flower field into two patches and programmed them to automatically change the spatial distribution of available nectar resources during the night. We performed our study with 36 individuals of male and female *G. mutica*. By using artificial flowers in a closed environment, we could track all flower visits and total nectar consumption of every individual in the group. Each individual carried an electronic ID tag and flowers were equipped with ID sensors. This also enabled us to detect and quantify a typical class of aggressive interactions between pairs of individuals directly at the artificial flowers fully automatically.

Our novel experimental set-up thus allowed us to perform a mostly automated experimental test of models of interference competition and resource defense with a mammal under the controlled conditions of the laboratory. This new approach may pave the way for further such studies with other groups of organisms.

## 2. Materials and methods

### (a) Subjects and housing

Experiments were conducted with 54 (36 females and 18 males) individuals of the small, (9-10g) neotropical nectarivorous bat species formerly identified as *Glossophaga soricina* (Pallas’s long-tongued bat). In view of the recent taxonomic revision of the *G. soricina* species complex (Calahorra-Oliart et al. 2021), it is relevant to note that the founders of our colony used in this and all our previous studies were caught at the Cueva de las Vegas, Municipio de Tenampulco, Mexico and transported to Germany in 1988 by Otto von Helversen. Thus they belong to the species *G. mutica*. Bats came from our captive colony and were older than one year as judged by finger joint ossification (Brunet-Rossini and Wilkinson 2009). They carried radio frequency identification (RFID) tags attached to cable tie collars (total weight of collar with tag = 0.2g, max. 2.4% of the body weight) that were removed after the experiment. Additionally, bats had numbered plastic split rings (A C Hughes Ltd., Middlesex, UK) around the forearm for visual identification. Temperature in the experimental and colony room was kept at 20-25°C, air humidity at 65-75%, and light conditions were 12:12 LD (light off at 16h).

### (b) Experimental set-up

In the experimental room ten artificial flowers with automated nectar delivery (Winter and Stich 2005) were mounted along a 4.2m bar at a height of 1.2m (Fig. 1). The distance between flowers was 0.4m. Flowers were divided into two groups of five to simulate two flower patches. Each patch was enclosed by a sheet-covered frame around the four sides and at the top to separate the groups of flowers spatially (Fig. 1). The only entrance to the patches was a 0.4m gap between the ground and the bottom end of this enclosure (Fig. 1, dashed line). From this entrance bats had to fly up vertically to reach the flowers, which increased the costs of moving between patches. A stepper-motor syringe pump delivered nectar via tubes and pinch valves to the artificial flowers. Nectar rewards were triggered by the interruption of an infrared light barrier at the flower opening. The RFID reader below the flower head identified a bat’s ID code. Flower visits (infrared light barrier interruptions) and ID sensor events were recorded during every experimental night. The reward schedule was configured using PhenoSoft Control (Phenosys GmbH, Berlin, Germany). Every detected event at a flower (including date, time, individual ID, duration of the event and amount of nectar delivered) was recorded for data analysis.

**Figure 1:**
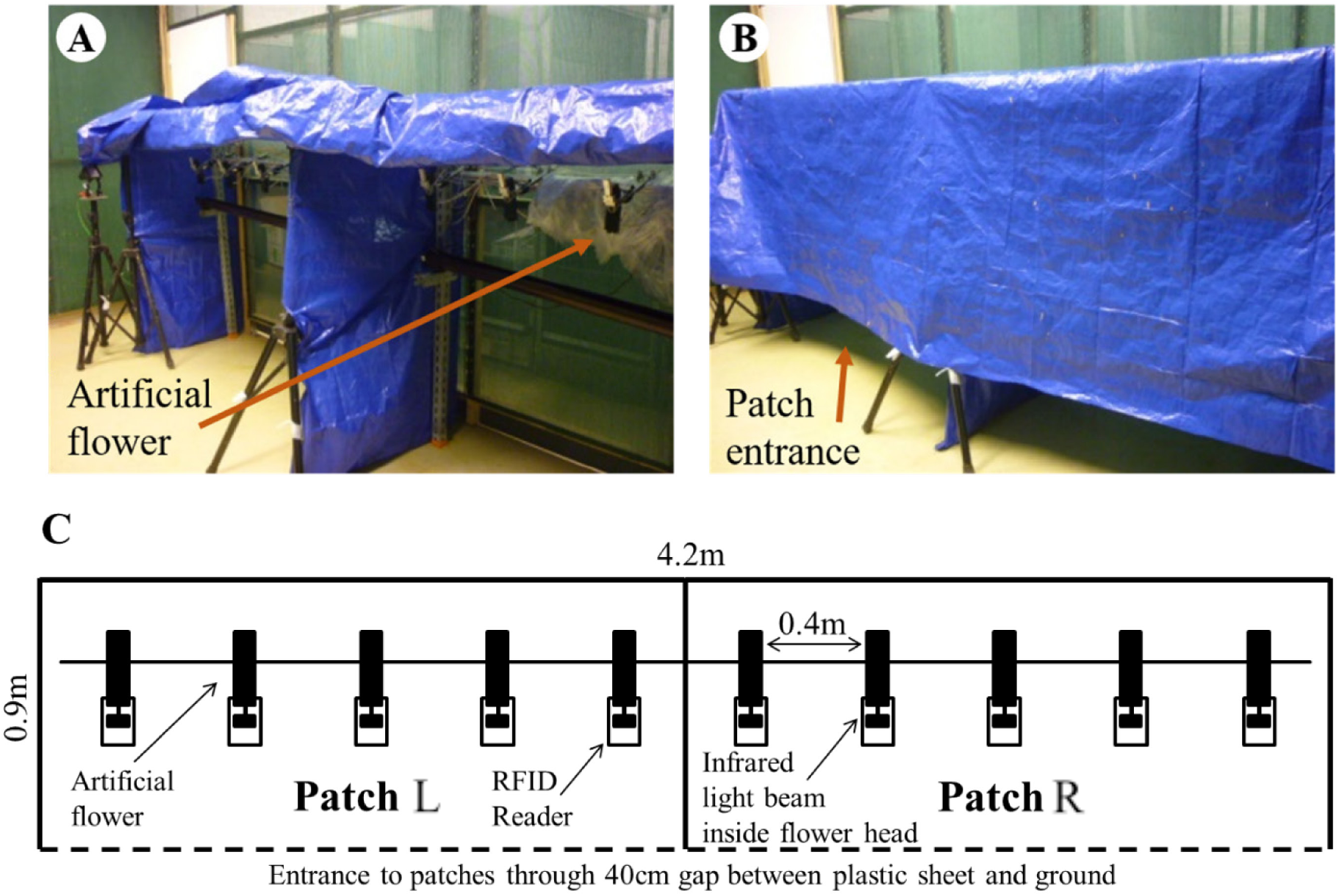
Experimental set-up consisting of two spatially separated patches of five flowers each. (A) The ten flowers were mounted 1.2m above ground. They were divided into two patches, L and R. (B) During experiments the patches were separated by plastic sheets. To make it more demanding for bats to enter a patch, the only entrance was through a 0.4m gap above the ground. (C) Schematic drawing of the experimental set-up from above. The dashed line indicates the side with the patch entrance.

### (c) Experimental procedure

Six bats were randomly selected from the colony (out of the total of ca. 150 individuals) and were tested simultaneously as a group. Four experimental groups consisted of three males together with three females (mixed groups), whereas one group consisted of six males, and four others of six females. No bats were re-used between experiments. We weighed all bats before the experiment.

During the nightly experiments, in addition to the nectar provided by artificial flowers, bats had access to pollen and water and to *6mL* of additional food containing 1.2g NektarPlus (Nekton, Keltern, Germany) and 1.8g milk powder (Milasan Folgemilch 2, Sunval Baby Food GmbH, Mannheim, Germany) dissolved in water. Rewards at flowers consisted always of *30μL* nectar (15% w/w sugar concentration, sucrose: fructose 1:2). Before the experimental schedule started, individuals were allowed to familiarize themselves with the set-up and the artificial flowers. Since during this training phase the plastic cover was removed, the two flower patches were not spatially separated and every flower visit was rewarded. This phase lasted for one to four nights until each bat visited the flowers regularly. The ID tag of one female of the first mixed group was not detected at any artificial flower during the first night and she was replaced by another female.

During the experiment, the two flower patches were covered and spatially separated (Fig. 1). Experimental nights were divided into two phases. During the first phase of the night only one of the two flower patches was rewarding, and therefore the resources were spatially clumped at a single location. The fixed time interval between rewards at each flower was 60s. During the second phase of the night both patches gave rewards, resources were evenly dispersed across the two patches, and the fixed time interval between two rewards at a flower was increased to 120s. Therefore, the amount of food available per unit time did not change during the whole night; only the spatial distribution of food changed from the clumped resource treatment with one patch rewarding (five flowers) during the first phase of the night to the dispersed resource treatment with two patches rewarding (ten flowers) during the second phase of the night. With this experimental schedule, the maximal amount of nectar the bats could collect was 108*mL*, which corresponds to *18mL* nectar per individual per night, roughly 150% of their daily requirement (Winter and von Helversen 2001). The side of the rewarding patch during the first phase of the night was chosen pseudo-randomly and the same patch was never chosen in more than two consecutive nights. For the mixed groups, the duration of the clumped resource treatment was six hours and the experiment lasted nine nights (seven nights for the first mixed group). For the same-sex groups, the duration of the first part of the night was variable (range = 4-8h, mean = 6h) and the experiment lasted eight nights for the male group and seven nights for the female groups.

### (d) Chasing behavior

We took the frequency of individuals chasing each other at the artificial flowers as an indicator of the intensity of aggressive interactions between group members. We developed a method to automatically detect and score chasing events using the computer-collected animal identification data from the RFID sensors and flower sensors. In a previous pilot study (Wintergerst 2018), three mixed groups of bats were video recorded for 24h over 14 nights, and the video data were synchronized to the computer-collected data. During this pilot study flowers were not covered by plastic sheets so that all flowers and the surrounding room were visible on video. From the analysis of the combined data we were able to identify the following pattern of visitation events that reliably indicated a chasing event between two identified individuals: *(i*) an identified bat visited a flower and (*ii*) its visit was instantaneously followed by the detection of a second bat, the chaser, that was detected very briefly (<200ms) and only by the ID sensor (detection range 5-7cm). Importantly, this second bat never attempted to drink and therefore did not insert its nose into the artificial flower and interrupt the light barrier inside the flower head. This distinguished such a chase from the occasional quick succession of two feeding visits by two bats at the same flower. This automated detection of chasing events not only saves considerable time for the experimenter, but also avoids human observer bias, a common drawback in video analysis. For the 24 hours of combined video data and automatically logged data, all 89 chasing events detected in the computer-logged data were confirmed by video (Wintergerst 2018). Therefore, we consider the algorithm for detecting chasing events in the logged data to be highly reliable. Of course, chases did not only occur at the artificial flowers. Thus, our chase numbers are only an indicator of chasing intensity between pairs of bats. For example, in one hour of video we observed 61 chasing events, but only five of those occurred during flower visits and were also automatically detected. However, since with our algorithm (see below) we detected a total of 2597 chasing events (35.8 ± 16.6 events per night during the experiment and only 4.1 ± 2.8 during the training nights, mean ± SD) for the 36 participating bats, we considered the automated approach adequate for quantifying within-group dominance relationships. The total number of individual detections per night constrains the number of chasing opportunities detectable with our method. Therefore, we calculated a *chase score* and a *chased score* by dividing the number of observed chases (directed to others or received from others, respectively) for each bat by the total number of detections for that bat on each night.

### (e) Statistical analysis

To investigate the difference in chasing behavior between males and females and between the resource treatments (one versus two rewarding patches) a Bayesian generalized linear mixed model (`MCMCglmm`, Hadfield 2010) with a binomial error structure and a parameter expanded prior was used. Body weight as an approximation of size and the full interaction between resource treatment and sex were included as fixed effects and the influence of these fixed effects on the proportion of chasing events was assessed. Experimental group and individual were included as random effects. The same model structure was used to address the question whether the proportion of visits on which the visitor was chased was influenced by these independent variables. If one or more individuals start to defend flowers and thus exclude others from drinking, nectar consumption should increasingly differ between individuals since the successful chaser should gain a higher nectar intake thus reducing the intake of the chased individuals. Therefore, the between-individual difference in nectar consumption over the course of the experiment was compared between experimental groups and resource treatments (clumped vs. dispersed). First, each individual’s total nectar consumption standardized by the number of hours of foraging during the clumped (one rewarding patch) and dispersed (two rewarding patches) resource treatment was determined for each experimental night. Then these data were used to calculate group standard deviations, separately for the males and females of each group. In order to assess the influence of resource defense on the individual differences in nectar consumption (standard deviation of nectar intake) we fit a `MCMCglmm` model with a Gaussian error structure and the following fixed effects: sex, experimental night (centered), and resource treatment (clumped or dispersed), as well as all two-way interactions. Again, we included group and individual as random effects.

By plotting individual nectar consumption during the last two nights of the experiment against the chase scores, two non-overlapping groups of males were detected, which were labeled dominant (territorial) and subordinate males, respectively. Such a clear pattern was not observed in females. The identification of dominant individuals was also supported by calculating the individual Glicko ratings (Glickman 1999; So et al. 2015) from all chasing events over the last two experimental nights in each group. In the Glicko rating algorithm individuals gain or lose ranking points based on their wins or losses and the rating of their opponent (Glickman 1999; So et al. 2015). Glicko ratings were analyzed using the `PlayerRatings` package in R (Stephenson and Sonas 2020). Based on nectar consumption, the frequency of chasing events and the individual Glicko group ranks (from 1 to 6, with 1 corresponding to the highest Glicko rating), each group contained individuals belonging to one of three different types of social status: female, dominant male, and subordinate male. To address the question whether nectar consumption varied depending on social status during the early and late stages of the experiment we used Welch’s tests and adjusted the p values using the Holms method for multiple comparisons.

All statistical analyses were conducted using R (Team 2021).

## 3. Results

### (a) Behavioral observations

The goal of our experiment was to investigate the sex-specific effects of resource defense in *Glossophaga mutica*, in addition to the potential influence of interference competition on individual nectar intake. Qualitative behavioral observations of four hours of video recordings revealed several behaviors that seem to be characteristic for some males, which according to further analyses (see below) we designated as dominant males. Instead of just visiting the flowers and leaving the patch as the other individuals did, dominant males remained hanging between the flowers within the patch for a significant amount of time (Fig. S1). When other individuals came close due to visits of directly adjacent flowers, dominant males often spread one wing in the direction of the other individual which could be interpreted as a threatening posture. Some individuals were attacked and chased away by dominant males while visiting artificial flowers. In this case, dominant males mostly attacked from above with their mouth wide open, and followed the intruder for a short distance. Sometimes the chasing escalated into fighting with both bats tumbling towards the ground and resuming their flight only shortly above the floor. In rare cases, these fights might have led to small injuries. One subordinate male had several fresh scratches on its wing that were not present before the experiment and that were possibly caused by bites (Fig. S2). After a successful flower defense, the dominant male normally visited most of the five flowers within the patch before returning to its hanging position between the flowers.

### (b) Example of nectar intake in one experimental group

One of the first striking observations we made was the uneven distribution of nectar consumed between the sexes and individuals. For example, in the first mixed group of bats tested, after only two nights the nectar consumption of two males was nearly reduced to zero, whereas the third male increased its consumption substantially (Fig. 2A). However, this pattern occurred mostly for males during the clumped resource treatment (Fig. @ref(fig: overview)). Nectar consumption of females was rarely as divergent as in males, especially in the mixed groups (Fig. @ref(fig: overview)). Later on the same nights when resources were dispersed over two patches, nectar consumption of males and females generally converged by the end of the experiment (Fig. 2B, Fig. @ref(fig: overview)).

**Figure 2:**
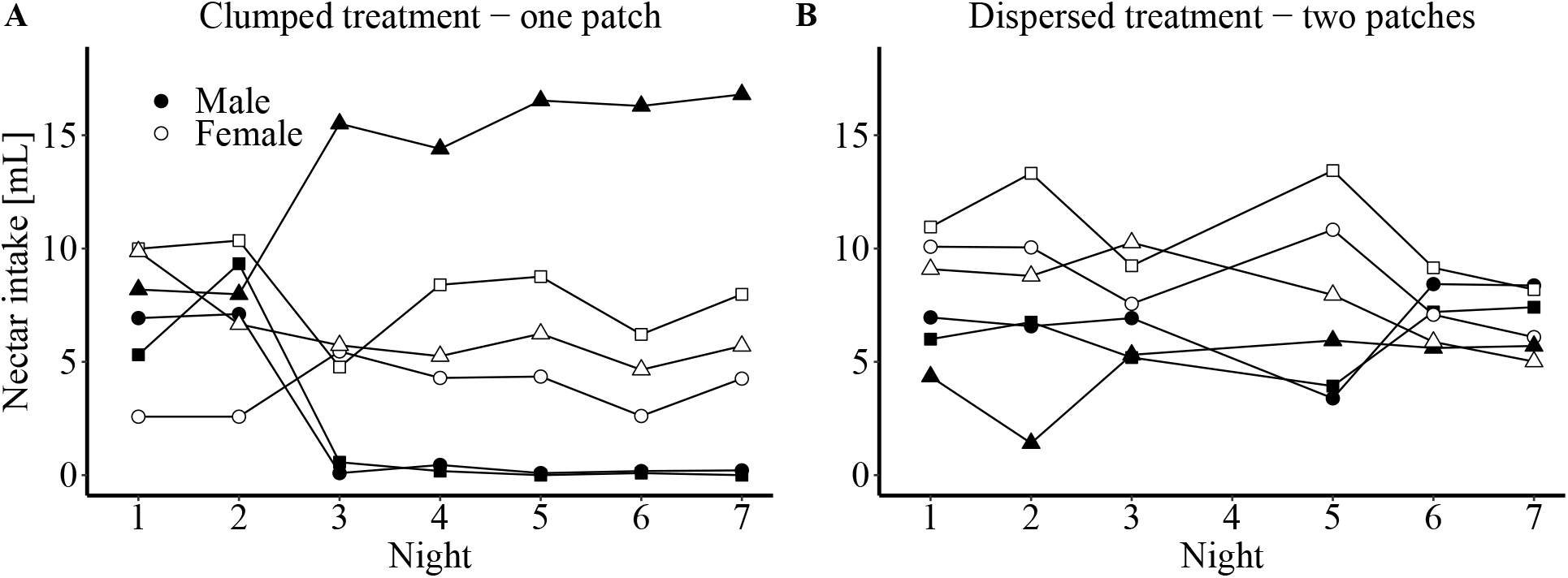
Example of the change of individual nectar consumption from the clumped treatment (A) to the dispersed treatment (B) during an experiment of one mixed group (3 males, 3 females, symbols show different individuals). (**A**). During the clumped resource treatment (first part of the experimental night) rewards were only available at one patch. The nectar consumption of two subordinate males approached zero after only two nights, whereas the third, dominant, male greatly increased nectar intake during the experiment (males filled symbols). Females (open symbols) on the other hand maintained a stable level of nectar intake. (**B**) During the dispersed resource treatment (second part of the experimental night) rewards were available at both patches. Under this treatment, individuals nearly equalized their level of nectar intake over the course of the experiment. The second part of night 4 was excluded due to technical problems.

### (c) Differences between sexes in frequency of chasing (chase score) and being chased (chased score)

In all mixed groups males chased other bats in front of flowers significantly more often than females did (Fig. 3A, Table 1). Notably, the frequency of females as active chasers in female-only groups was higher than chasing by females in the mixed groups (Fig. 3A). Although the rate of nectar availability remained constant throughout the night and only the spatial distribution of the resources changed, chase frequencies were significantly lower during the dispersed resource treatment when rewards were available at both patches (Table 1). There was no significant difference between the sexes in how often a bat was chased by another individual (Fig. 3B) but individuals were chased less during the dispersed resource treatment (Table 1). Weight as an indicator of size had no significant effect on either the chase score or the chased score (Table 1).

**Figure 3:**
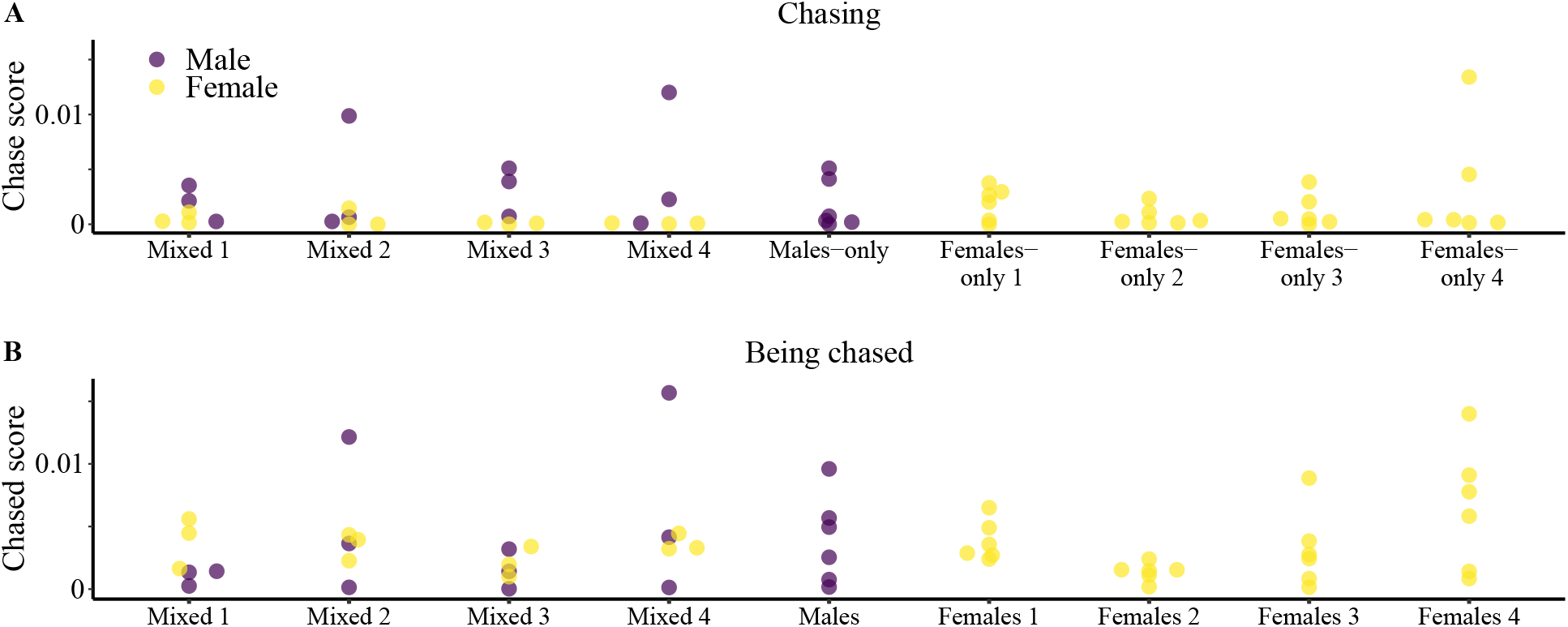
Sexes differed in the frequency of chasing or being chased during the clumped resource treatment. (**A**) Males (dark symbols) chased others significantly more than females did (light symbols, Table 1). Shown are the individual proportions of chasing events (chase scores) over the whole experiment. Notably, in the females-only groups some females chased more than any female in the mixed groups. (**B**) Being chased by other bats did not differ significantly between sexes (Table 1).

**Table 1:** Summary of fixed effects from generalized linear mixed-effects models of chasing frequency and the frequency of being chased.

| Model | term | estimate 95% credible interval | pMCMC |
| --- | --- | --- | --- |
| Chase score | (Intercept) | -5.20 (-11.54, 0.64) | 0.0970 |
|  | sex (female) | **-1.33 (-2.53, -0.2)** | **0.0290** |
|  | phase | **-0.46 (-0.76, -0.17)** | **0.0050** |
|  | weight | -0.11 (-0.75, 0.5) | 0.7470 |
|  | sex (female):phase | -0.06 (-0.46, 0.34) | 0.7490 |
| Chased score | (Intercept) | **-5.24 (-10.06, -0.41)** | **0.0300** |
|  | sex (female) | 0.72 (-0.12, 1.62) | 0.1060 |
|  | phase | **-0.95 (-1.22, -0.67)** | **0.0005** |
|  | weight | -0.07 (-0.58, 0.41) | 0.7810 |
|  | sex (female):phase | -0.11 (-0.44, 0.21) | 0.5340 |

Note: Fixed estimates whose credible intervals do not span zero are shown in bold. pMCMC = posterior probability

### (d) Differences in nectar intake over time and between sexes and treatments

Resource defense should lead to a larger between-individual difference in nectar consumption (Brown 1964). Differences in nectar consumption were quantified as the standard deviation of nectar intake in each group, separately for males and females. During the clumped resource treatment, the standard deviation was higher for males than for females (Table 2, Fig. S24) and increased significantly over time for males but not for females (Table 2, Fig. S24). For females in the clumped resource treatment the increase in standard deviation was significantly smaller than in males (significant interaction between sex and night, Table 2), and was not itself significant (estimate = 0.02, 95% CI = 0, 0.06). Compared to the clumped resource treatment, in the dispersed resource treatment the effect of experimental night was significantly lower for males (interaction between treatment and night, Table 2), but not for females (estimate = 0, 95% CI = −0.02, 0.02). Moreover, in the dispersed resource treatment there was no significant increase in standard deviation over the course of the experiment in males (estimate = 0.02, 95% CI = −0.01, 0.05) nor in females (estimate = 0.03, 95% CI = 0, 0.06). Overall, for both males (significant effect of treatment) and females (estimate = −0.11, 95% CI = −0.16, −0.06) the standard deviations were generally higher in the clumped than in the dispersed resource treatments.

**Table 2:**
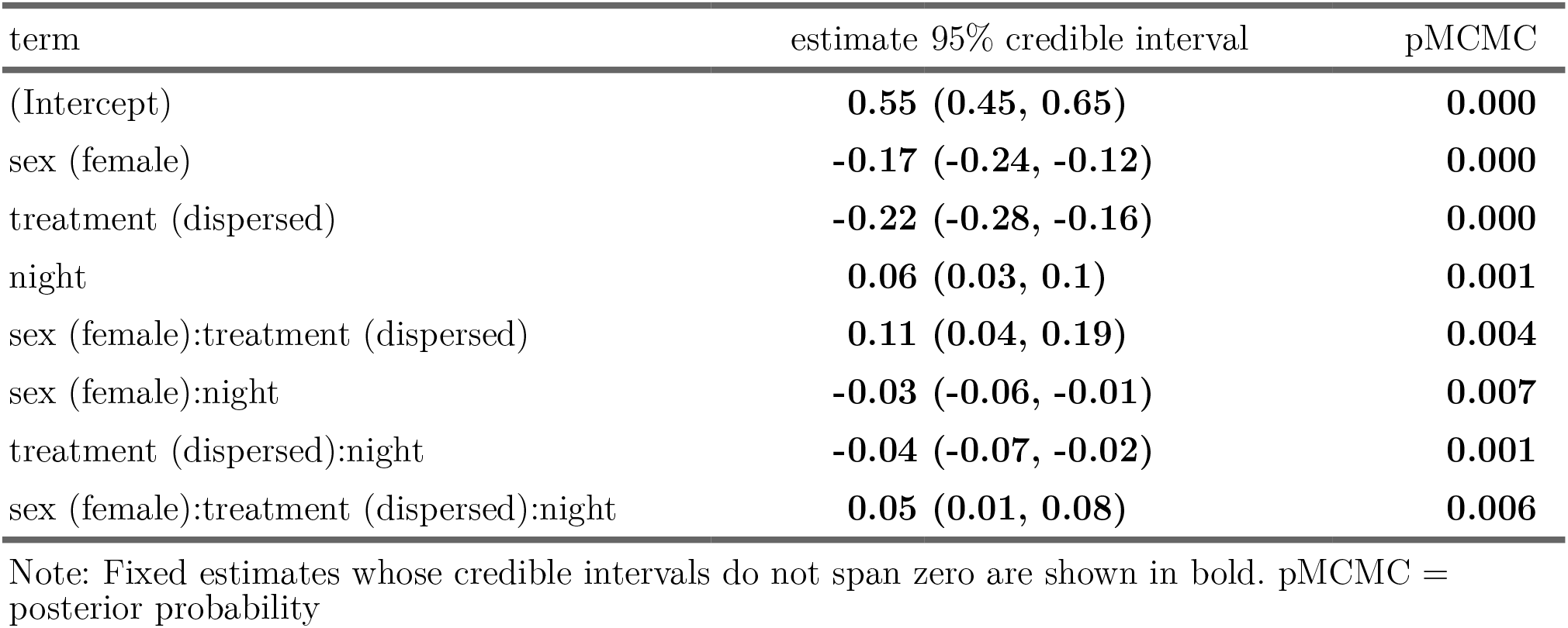
Summary of fixed effects from a generalized linear mixed-effects model of the standard deviation of nectar intake over time.

| term | estimate 95% credible interval | pMCMC |
| --- | --- | --- |
| (Intercept) | **0.55 (0.45, 0.65)** | **0.000** |
| sex (female) | **-0.17 (-0.24, -0.12)** | **0.000** |
| treatment (dispersed) | **-0.22 (-0.28, -0.16)** | **0.000** |
| night | **0.06 (0.03, 0.1)** | **0.001** |
| sex (female):treatment (dispersed) | **0.11 (0.04, 0.19)** | **0.004** |
| sex (female):night | **-0.03 (-0.06, -0.01)** | **0.007** |
| treatment (dispersed):night | **-0.04 (-0.07, -0.02)** | **0.001** |
| sex (female):treatment (dispersed):night | **0.05 (0.01, 0.08)** | **0.006** |

Note: Fixed estimates whose credible intervals do not span zero are shown in bold. pMCMC = posterior probability

### (e) Social status and its effects on nectar intake

When plotting chasing events against nectar consumption the data for males fell into two non-overlapping groups. The males of one cluster (Fig. 4A, inside dashed oval) chased other individuals and consumed more nectar than the other males. This cluster included only one male from each of the four mixed groups but two males from the single males-only group. These six males were categorized as “dominant”. The second cluster of males (Fig. 4A, outside and below dashed oval) was characterized by a low frequency of chasing and low nectar consumption. These males were categorized as “subordinate”. In females such a pattern did not emerge (Fig. 4B). This classification was also supported by the highest social dominance scores as estimated by Glicko ratings in each group (Figs. S25, S26) and the observation that there was generally an inverse relationship between the frequency of chasing and the frequency of being chased (Fig. S27). While in the females-only groups some females chased other females more frequently, only three of these females (one in group 1 and two in group 4) would be classified as dominant using the same cut-off criteria we used for the males (Fig. 4B). While in females-only group 4 the two females were the individuals with the highest Glicko ratings, in females-only group 1 the female with the highest nectar consumption and chase score did not have the highest Glicko rating (Figs. S25, S26). During the last two nights of the experiment in the clumped resource treatment, the highest nectar intake was observed in dominant males, with an intermediate intake in females, and lowest nectar intake in subordinate males (Fig. 5). In contrast, in the dispersed resource treatment there were no detectable differences between the nectar intake of dominant and subordinate males at any stage of the experiment (Fig. 5), while the subordinate males had a significantly lower nectar intake than females in the first two, but not in the last two experimental nights (Fig. 5). Finally, the subordinate males increased their nectar intake from the clumped to the dispersed treatment, but the difference was only significant for the last two experimental nights (Fig. 5). While there was a correspondent decrease in the nectar intake of dominant males, it was not significant, most likely due to the small sample size (n = 6, Fig. 5). In females there were no significant differences in nectar intake from the clumped to the dispersed resource treatment at any stage of the experiment (Fig. 5)

**Figure 4:**
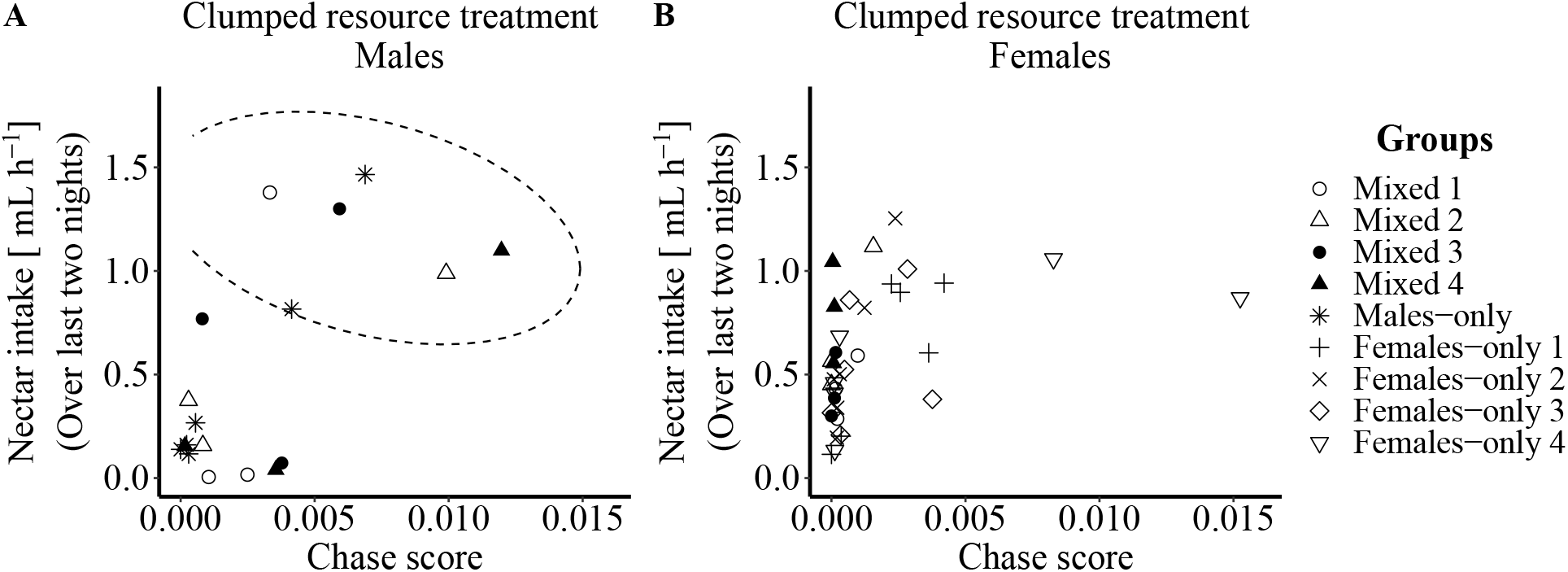
Influence of chasing frequency (chase score) on nectar intake in the clumped resource treatment during the last two nights of the experiment. (**A**) Males that more often chased other males also consumed more nectar. Males were divided into two non-overlapping groups by considering the chasing frequency and the amount of nectar an individual received during the clumped resource treatment at the end of the experiment. Dominant males (inside dashed line oval) met two criteria: they chased other individuals at flowers more frequently (>0.003) and received more nectar (>0.75*mL h*^-1^) during the clumped resource treatment. Individuals outside the dashed line oval were categorized as subordinate males. (**B**) Nectar consumption of females did not generally depend on the chase score during the clumped resource treatment and non-overlapping groups did not emerge.

**Figure 5:**
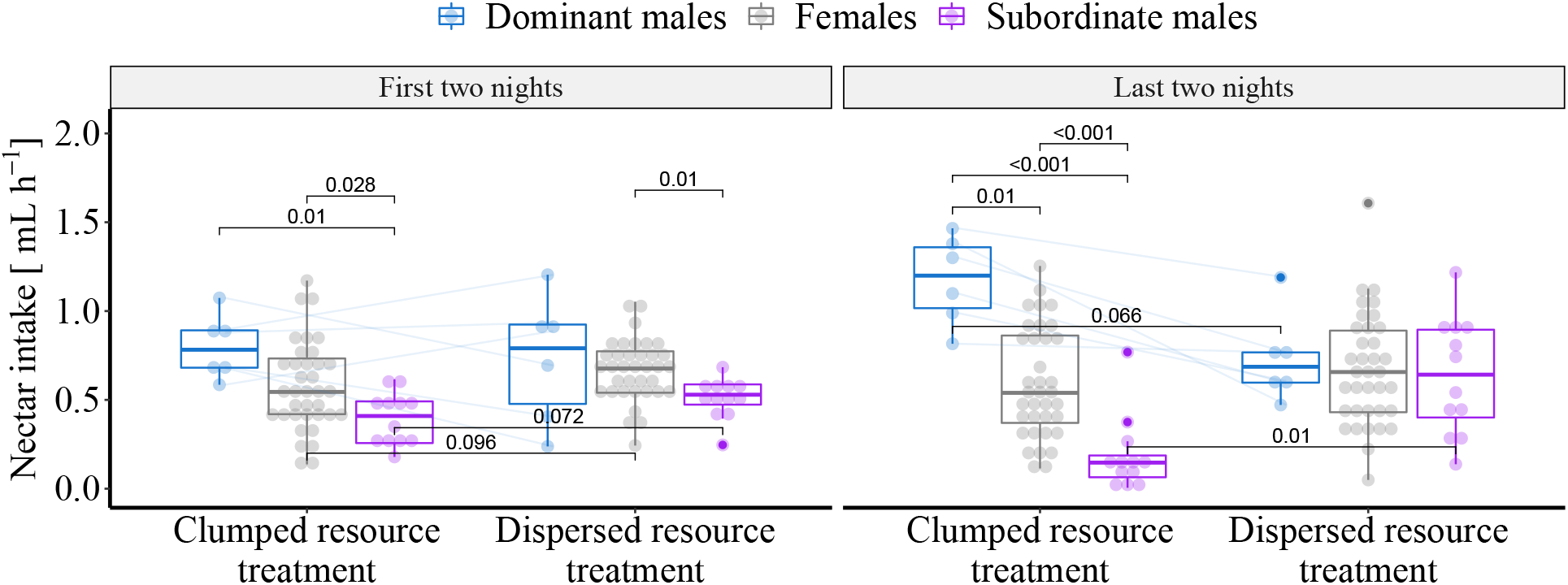
Comparison of nectar intake during the first and last two nights of the experiment depending on sex and social status. During the clumped resource distribution (left in each panel), already at the beginning of the experiment (left panel) subordinate males collected significantly less nectar than dominant males and females. At the end of the experiment (right panel), females, dominant and subordinate males differed in their nectar consumption, but only during the clumped and not during the dispersed treatment. During the dispersed resource treatment at the beginning of the experiment subordinate males received less nectar than females, but this difference disappeared by the end of the experiment. Numbers above brackets are the p values from unequal variance T tests (Welch’s tests), adjusted for multiple comparisons using the Holms method. Contrasts between treatments were from paired Welch’s tests. For clarity, only p values smaller than 0.1 are shown. Each status group represents data from the same individuals. For clarity, only the lines connecting the same dominant males are shown.

## 4. Discussion

Similar to observations in free-living *Glossophaga* populations (Lemke 1984; Tschapka 2003), in this experiment *G. mutica* competed for nectar not only by exploitation but also by interference competition. However, the results show that the predisposition to defend resources and the influence of interference competition on individual nectar intake differed significantly between the sexes. Only a subset of individuals, most notably males in the mixed-sex groups, successfully defended flower patches. Dominant individuals were characterized by the highest frequency of chasing other individuals away from profitable flowers (chase scores), by the highest Glicko ratings, and by a substantial increase in nectar intake during the time periods of active defense by the end of the experimental run. Although the dominant males in the mixed groups chased females and other males about equally often, only the nectar intake of subordinate males but not of the females was affected by this behavior. Thus, male-initiated interference competition increased the difference in nectar intake between males but did not affect females. The frequency of aggressive interactions was higher, and males only defended resources successfully when the available nectar was concentrated at only one flower patch. This supports the hypothesis that clumped resources favor an increase in aggressive interactions (Grant 1993).

### (a) Sex-dependent resource defense and its differential effect on nectar intake, depending on social status

To our knowledge, this study is the first report of sex-dependent differences in resource defense behavior of neotropical nectar-feeding bats. In mixed sex groups, females seemed to be much less affected by the behavior of dominant males whereas subordinate males were excluded at least partially from the defended flower patch. This finding is consistent with observations of free-flying *G. commissarisi*, in which males visited on average a smaller number of artificial flowers than females did (Nachev and Winter 2019), presumably because of interactions with other males. There are two possible explanations for this differential effect on subordinate males and females. On the one hand, dominant males might just not be capable of excluding females. On the other hand, dominant males could tolerate females in their defended patch because they might receive additional benefits, for example tolerating females could lead to an increase in (future) mating opportunities. Similar social dynamics have been described in the insectivorous bat species *Myotis daubentonii* (Senior et al. 2005). Dominant males of this species temporarily exclude other males from profitable habitats whereas females are tolerated and in addition to securing access to resources, the successful exclusion of other males has been shown to increase the reproductive success of dominant males (Senior et al. 2005). Similarly, it has been observed that male purple-throated carib hummingbirds *(Eulampis jugularis)*, which successfully defend highly profitable feeding-territories against other males while sharing the available resources with females, experienced an increase in their mating success (Temeles and Kress 2010).

However, in our experiment dominant males chased females about as often or slightly more often as they chased subordinate males (Table 1, Fig. S22). If females were able to feed in the defended patch because dominant males tolerated them due to potential additional benefits, it could be that the detected chasing events by dominant males differed in quality depending on the sex of the intruder. This was not further quantified in the current study but could potentially be investigated using audio recordings (Knörnschild et al. 2010; Hörmann et al. 2020). We extracted the frequency of chasing events from data automatically recorded at artificial flowers (successive detection of two different IDs while and after the first was feeding at the flower). Therefore, it was not possible to determine if males showed behavioral differences when chasing other males in comparison to chasing females. The recorded video revealed that individuals chased each other not only directly at the artificial flowers but also in other areas of the flower patch. Since individuals could only be identified by their ID tags directly at the ID reader attached to artificial flowers the sex of individuals chasing each other in other areas of the experimental room remained unknown. However, after the experiment some subordinate individuals showed marks from small injuries at their wings (see example in Fig. S2) and such marks were only observed in males. This could be an indication that dominant males directed more aggression (biting) towards subordinate males than towards females. Such sexual dimorphism in aggressive resource defense is also known from other nectar-feeding vertebrates, like hummingbirds. The beaks of the males of some territorial hummingbirds seem to be specifically adapted as intrasexually selected weapons (Rico-Guevara et al. 2019).

### (b) Some observations from the single-sex groups

Generally, females showed lower chasing frequencies, but, surprisingly, some females in the females-only group showed an increased nectar consumption and chasing frequency, compared to the females in the mixed groups (Figs. 4B, S25, S26). This observation prompted us to contrast the behavior of females in the two different group types (single-sex versus mixed-sex). Females in the single-sex groups had higher chase scores compared to females in mixed groups (Table S1). In single-sex groups, but not in mixed groups there was a higher frequency of chases in the clumped than in the dispersed resource treatment (Table S1). The chased score was only affected by treatment but not by group type and was higher in the clumped than in the dispersed resource treatment (Table S1). Over the course of the experiment, the standard deviations in nectar intake increased for females in the single-sex, but not in the mixed groups (Table S2). This increase was only significant in the clumped, but not in the dispersed resource treatment (Table S2). The standard deviations were higher in the single-sex groups than in the mixed groups, both in the clumped (group type effect in Table S2) and in the dispersed resource treatments (estimate = −0.06, 95% CI = −0.12, −0.01). Thus, it appears that in the absence of male individuals, some females exerted dominant behavior over the other females, similar to males. These findings are similar to the social structure of resource defense found in some nectar-feeding bird species. For example, in free-living ruby-throated hummingbirds females also have lower levels of defense (Rousseu et al. 2014). Moreover, although both male and female *Eulampis jugolaris* hummingbirds defend feeding territories during the non-breeding season, males are always dominant over females (Wolf and Hainsworth 1971; Temeles et al. 2005). It would be interesting to better understand why females were less affected by the aggressive resource defense behavior of dominant males compared to subordinate males and why females themselves did not consistently monopolize the profitable patch against other females, not even in the females-only groups. One possibility is that females do not need to defend flowers when a dominant male is already reducing the number of flower visitors and thus increasing the amount of food available.

In all mixed sex groups, only one male per group became dominant and successfully defended flowers, whereas in the males-only group two males exhibited dominant behavior (Fig. 4A). A closer look at the nectar consumption at each flower revealed that on the last night of the experiment these two males had nearly monopolized different flowers within the same patch rather than sharing access to the same flowers (Fig. S28). Such flower or patch partitioning was also observed in the females-only groups (Figs. S33–S36), but rarely seen in the mixed groups (Figs. S29–S32). The successful resource defense by two individuals in the male-only group showed that resource defense can occur independent of the presence of females, but, this was only based on a single observation.

### (c) Social status and social hierarchy

Although the position of the rewarding patch during the clumped resource treatment changed between the nights between the left and right, usually the same male continued to successfully defend the patch, especially in the mixed-sex groups (Figs. S28–S32). This means that males defended the resources themselves and not a particular location. Furthermore, this shows that even after changing the location of the defended patch the same individuals were usually able to succeed in re-establishing their dominance against other males, indicating a stable hierarchy at least for the duration of the experiment. The flower utilization pattern in females-only groups was not as consistent (Figs. S33–S36).

The ability of an individual to successfully defend and monopolize resources is often correlated with distinct physical characteristics such as body size (Searcy 1979). However, in our results weight as an approximation of size did not correlate significantly with the chase score of individuals (Table 1) and therefore did not predict which male succeeded in defending a flower patch. Another factor that could influence the success in defending flowers is age and therefore experience (Yasukawa 1979; Arcese 1987). Since we could only discriminate between young and adult animals, we cannot dismiss age and experience as a predictor of successful flower defense.

In this study, subordinate males received considerably less nectar than dominant males and females (Fig. 5). However, except in mixed group 1, subordinate males were rarely completely excluded from the flower patch and their average nectar intake during the clumped resource treatment was still 0.3 ±0.18 *mL h*^-1^ (mean ±SD). This result is in accordance with observations of free-living *G. soricina* in Colombia. There, subordinate bats exploited the flowers defended by other individuals as soon as the dominant bat temporarily ceased defending (Lemke 1984). Furthermore, in our study the frequency of chasing events decreased significantly during the dispersed resource treatment in the second part of the night (Table 1). This supports the theoretical prediction that aggressive defense behavior increases when resources are spatially concentrated (Grant and Guha 1993). However, since the sequence of treatments was not controlled in this experiment, other factors (e.g., time) cannot be ruled out. With the current data we cannot answer whether the dominant males would successfully defend a patch if the treatment changed from dispersed to clumped, but we believe this is a different question that should be addressed separately. Resource defense should only occur when the energy gain outweighs the cost of aggressive interactions (Brown 1964). Thus, our results could be explained by the decrease in quality of the defended patch once its nectar supply rate dropped to half. This is also supported by the very low number of chases observed during training when the flowers gave unrestricted rewards and were not separated in discrete patches. Together, these results suggest that along the different degrees of territorial behavior, resource defense observed in *Glossophaga* seems to represent a transient monopolization of resources instead of a longer-term permanent exclusion of intruders.

### (d) Conclusion

Although flower defense behavior of *G. mutica* was investigated in a laboratory setting, we observed similar behavior as described in free-living *Glossophaga* populations. Our results revealed a sexual dimorphism in flower defense behavior in mixed-sex groups. Only males successfully defended flower patches and excluded other males from their defended resource, whereas females remained unaffected by this male behavior and continued to visit the flowers guarded by a male. In the absence of males females also defended flowers against other females, but not as consistently as males. This observed pattern is similar to resource defense behavior observed in other nectar-feeding vertebrates. Furthermore, we could show that the frequency of aggressive interactions was, as predicted, higher when resources were clumped in one patch and transient. Future studies with free-living populations have to be conducted to assess how frequent and important resource defense in these nectar-feeding bats is and if males that are successful in defending resources have additional fitness advantages.

## Supplementary material

### Video analysis

There were 89 chase occurrences observed (f->f 4 times, f->m 2 times, m->f 59 times, m->m 24 times). Every time the algorithm marked an event as a chase event, there were two individuals following each other. Some chase sequences did not get detected. The individual that chased never drank immediately after the chase at the same flower location where the chase occurred. There were 16 incidences that were difficult to classify by observation or did not appear to be aggressive interactions.

f->f appear to be less aggressive
f->m appear aggressive
m->f appear aggressive
m->m appear aggressive

## Supplementary figures

**Figure S1:**
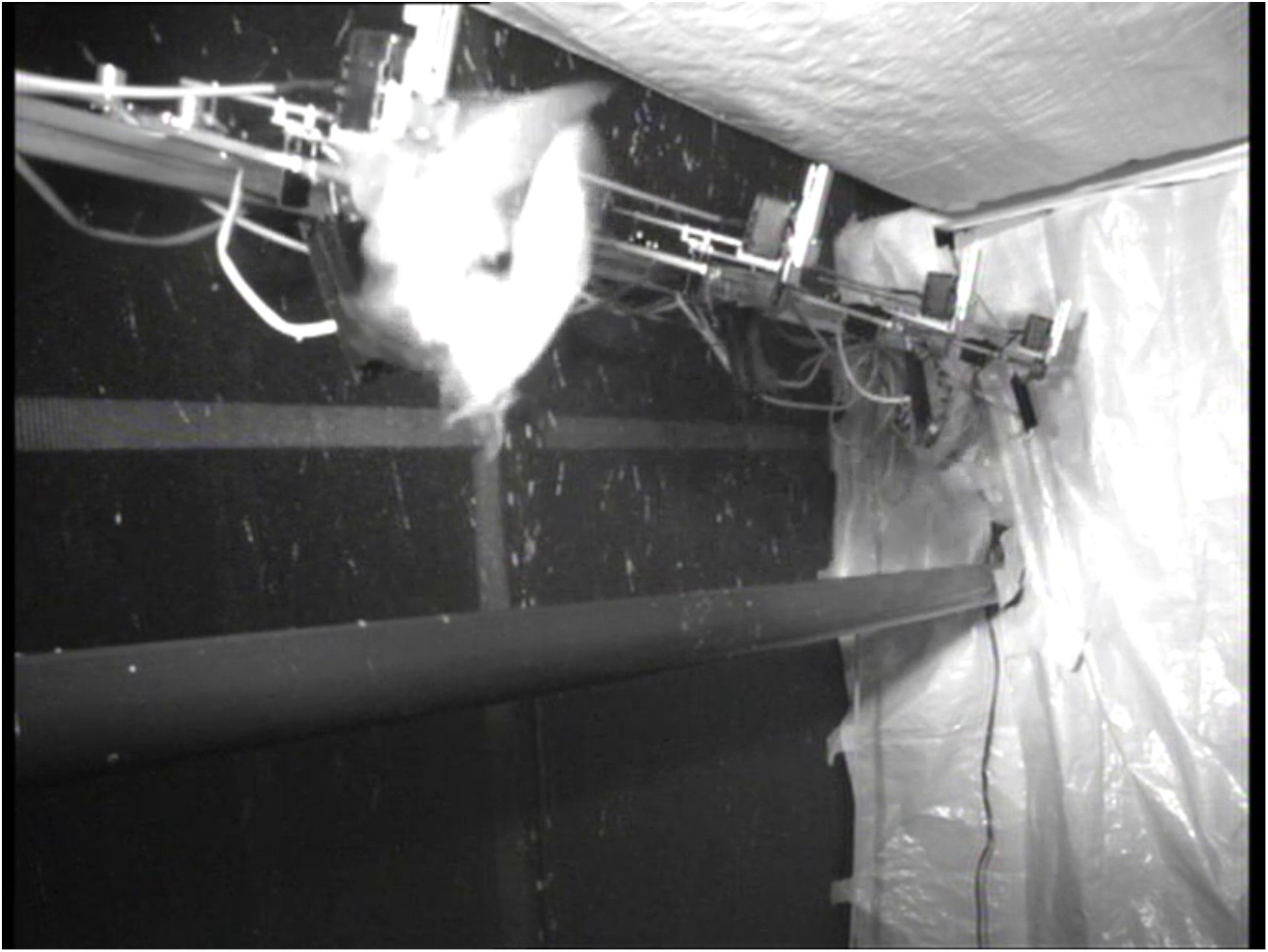
Video of the dominant male in mixed group 3 chasing all bats approaching the rewarding flowers in the rewarding patch during the clumped resource treatment.

**Figure S2:**
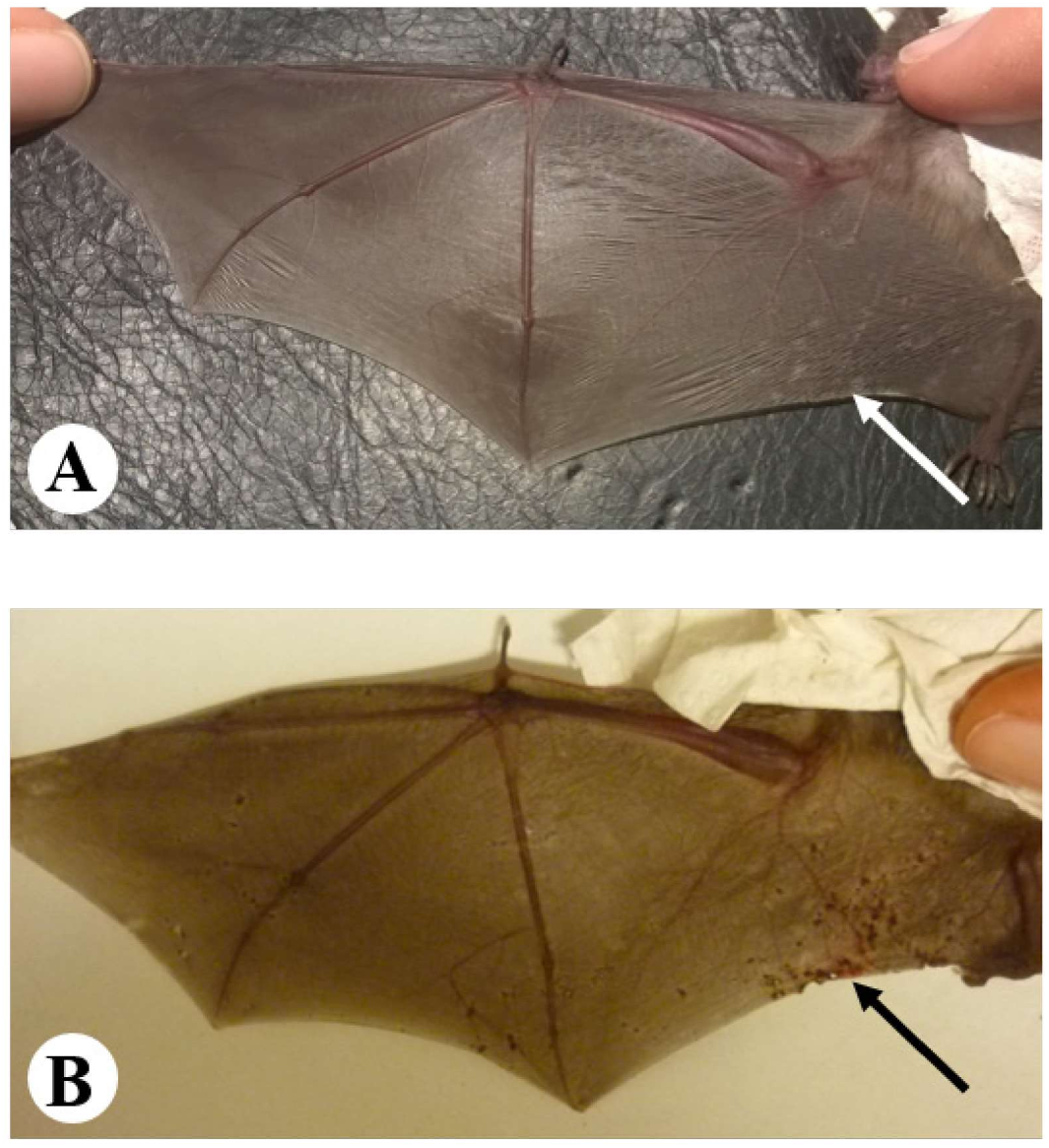
Wing images of a subordinate male from mixed group 4. The same individual was photographed before (**A**) and after the experiment (**B**). The black arrow points to the scarred location due to wing injuries, purportedly caused by the dominant male.

**Figure S3:**
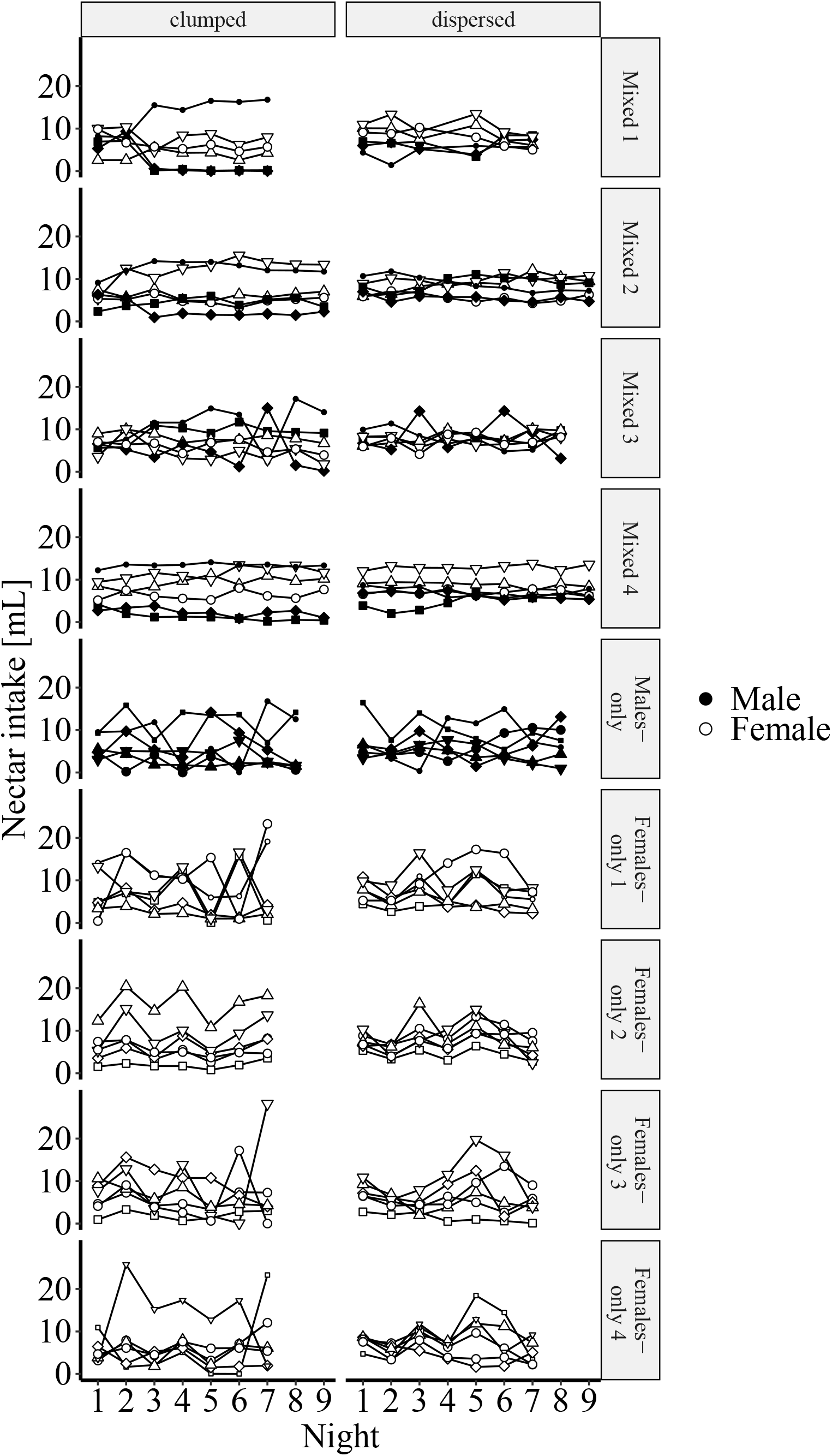
Change of individual nectar consumption from the clumped treatment (left) to the dispersed treatment (right) for all experimental groups (rows). Males are shown with filled symbols and females with open symbols. Small symbols indicate dominant individuals, according to proportion of chasing and nectar intake during the last two nights of the experiment.

**Figure S4:**
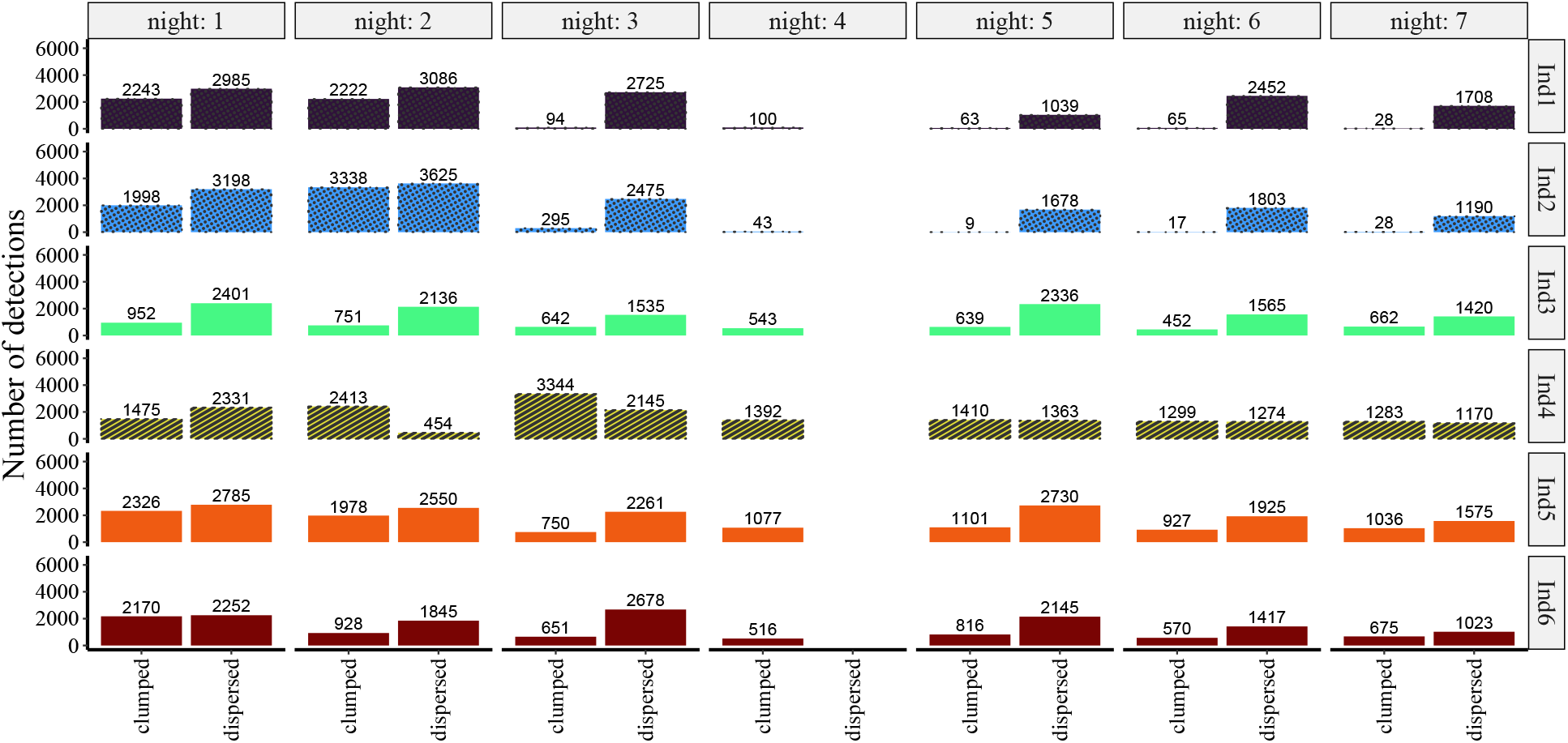
Raw number of detections for all bats in mixed group 1. Dotted bars represent subordinate males, striped bars represent dominant males and unhatched bars - females.

**Figure S5:**
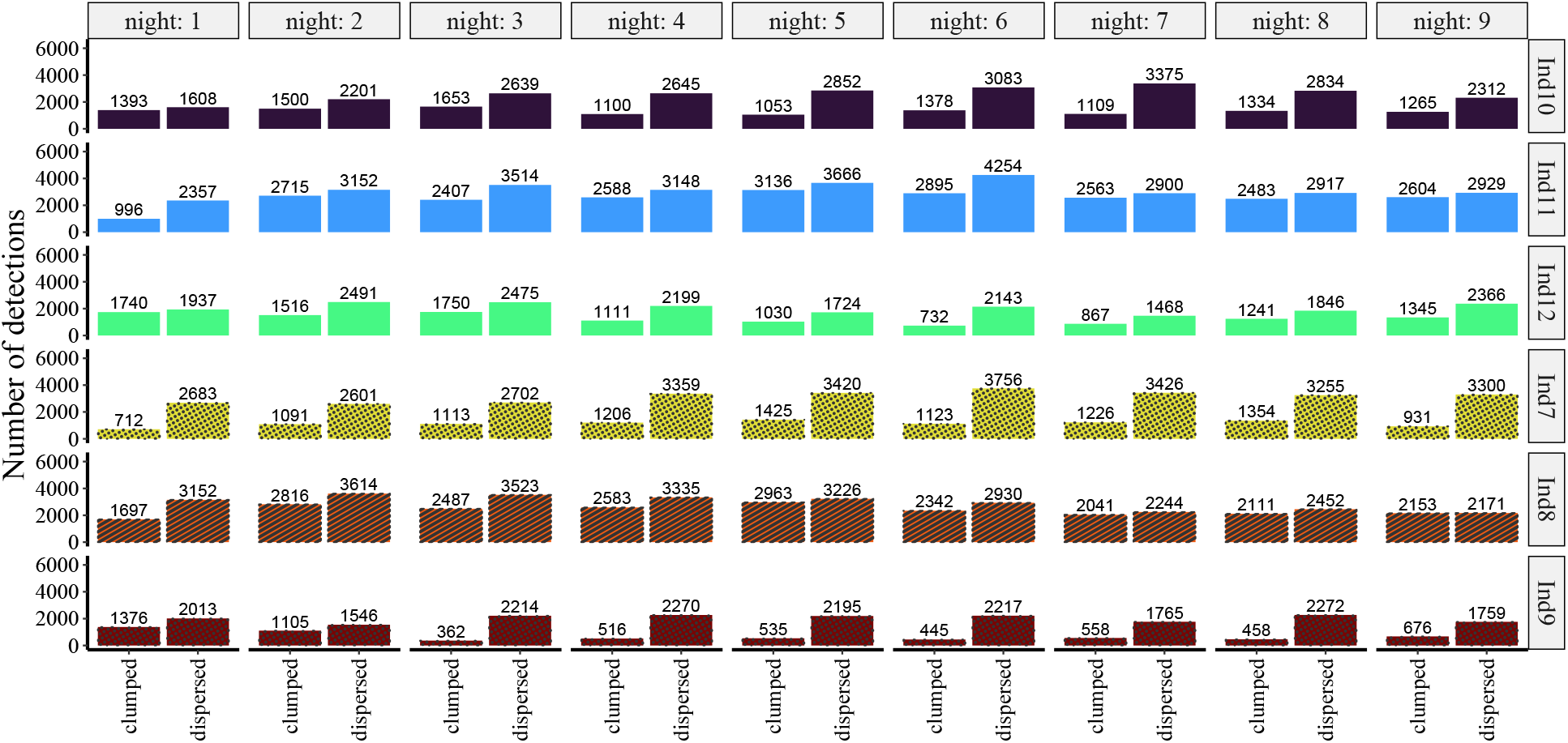
Raw number of detections for all bats in mixed group 2. Same notation as in Fig. S4, but the colors correspond to different individuals.

**Figure S6:**
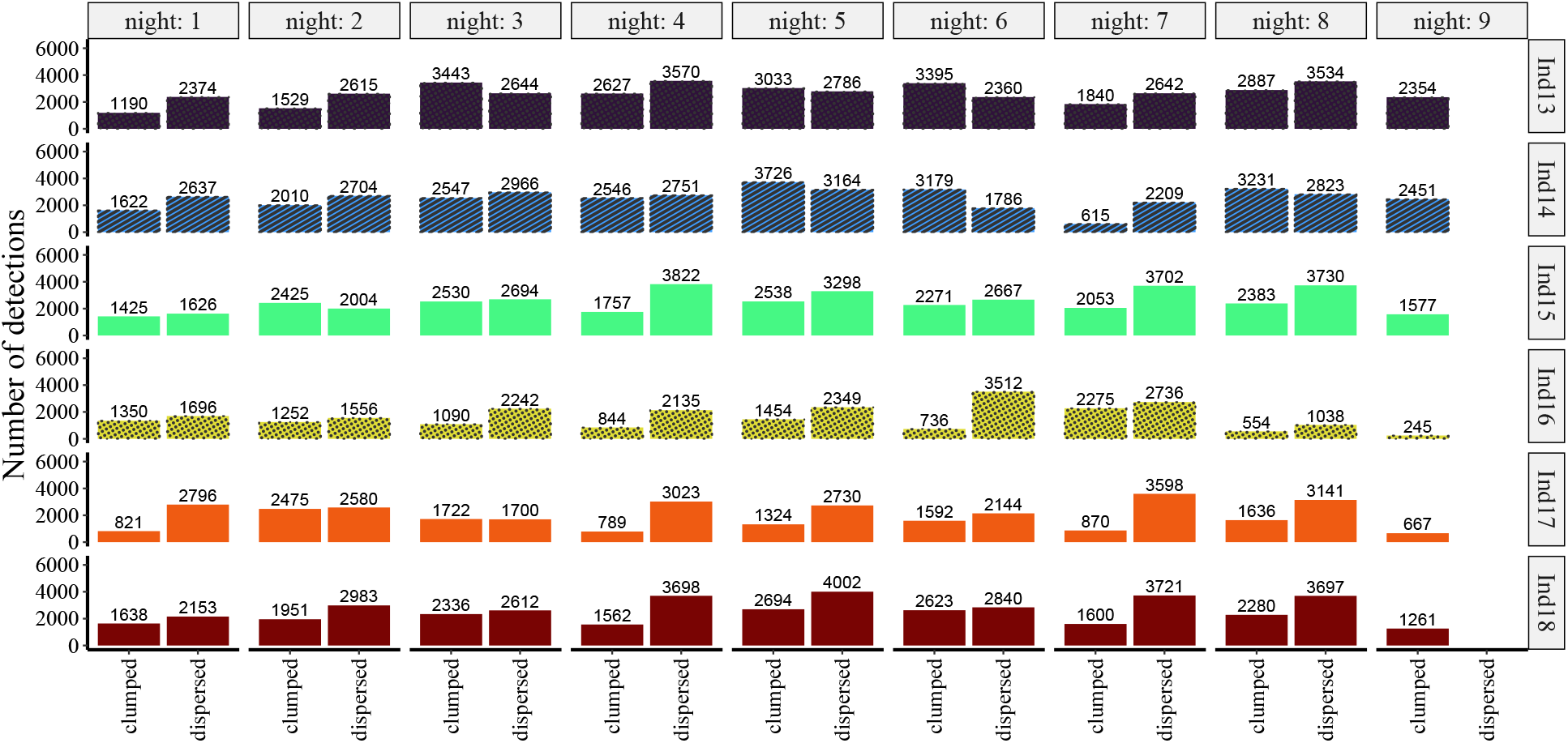
Raw number of detections for all bats in mixed group 3. Same notation as in Fig. S4, but the colors correspond to different individuals.

**Figure S7:**
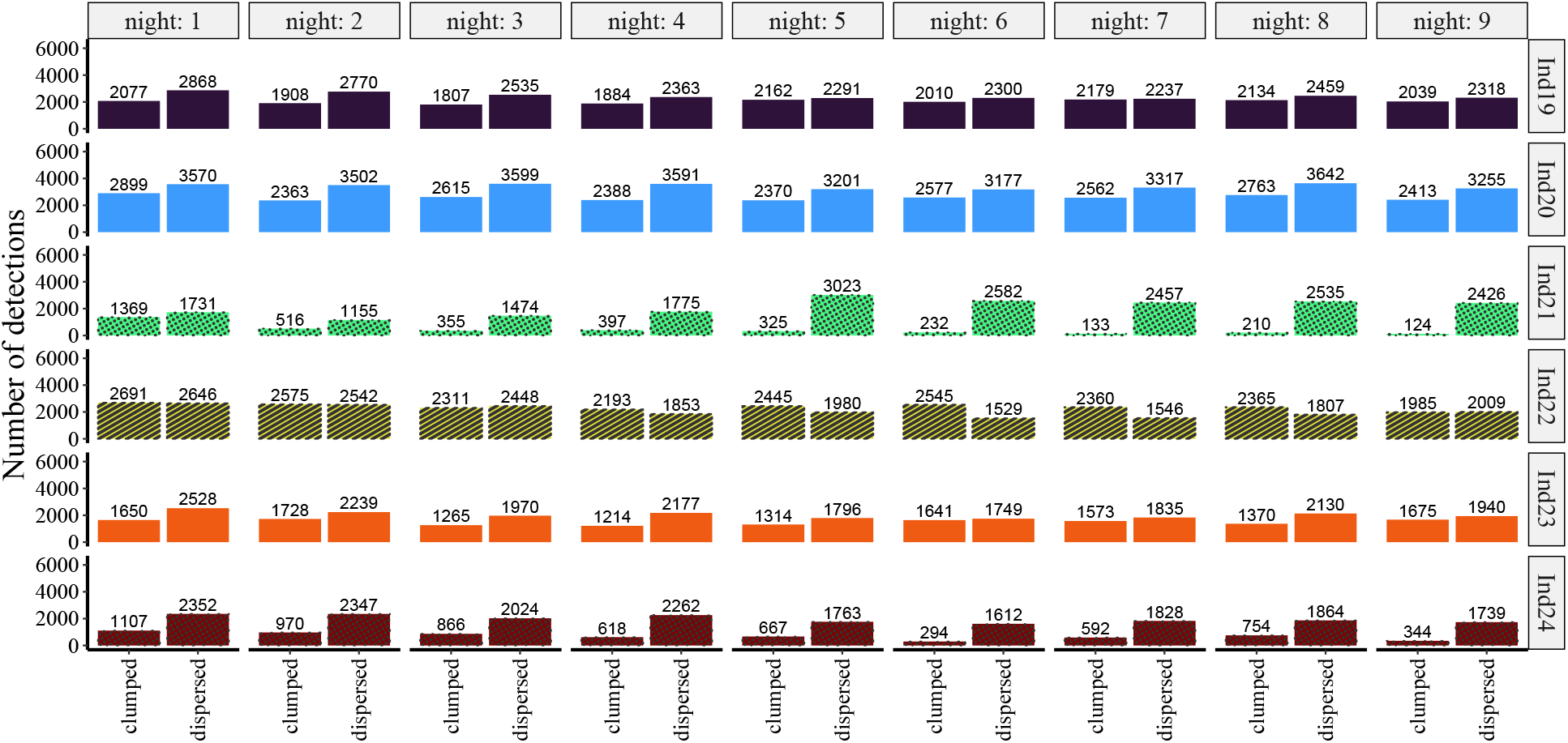
Raw number of detections for all bats in mixed group 4. Same notation as in Fig. S4, but the colors correspond to different individuals.

**Figure S8:**
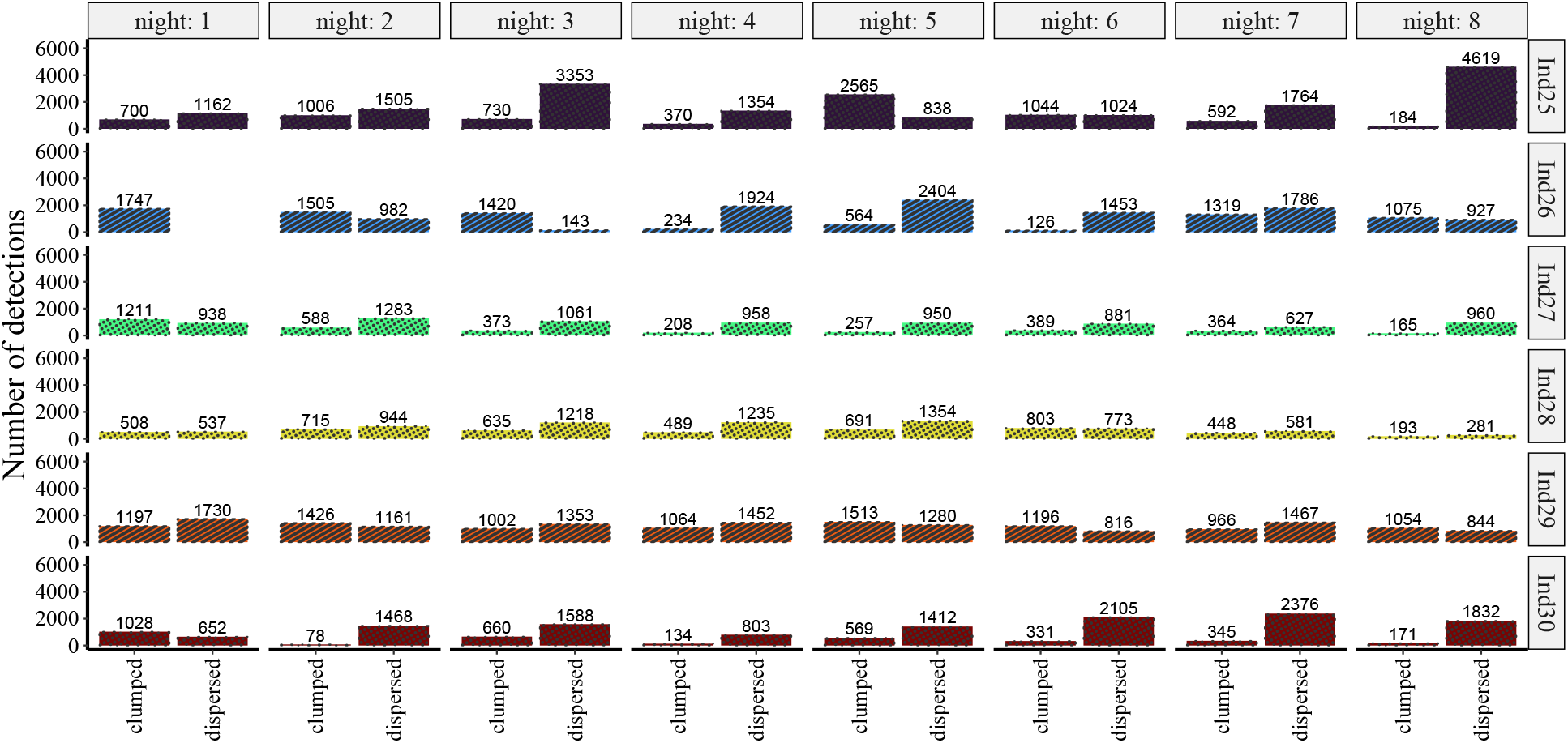
Raw number of detections for all bats in males-only group. Same notation as in Fig. S4, but the colors correspond to different individuals.

**Figure S9:**
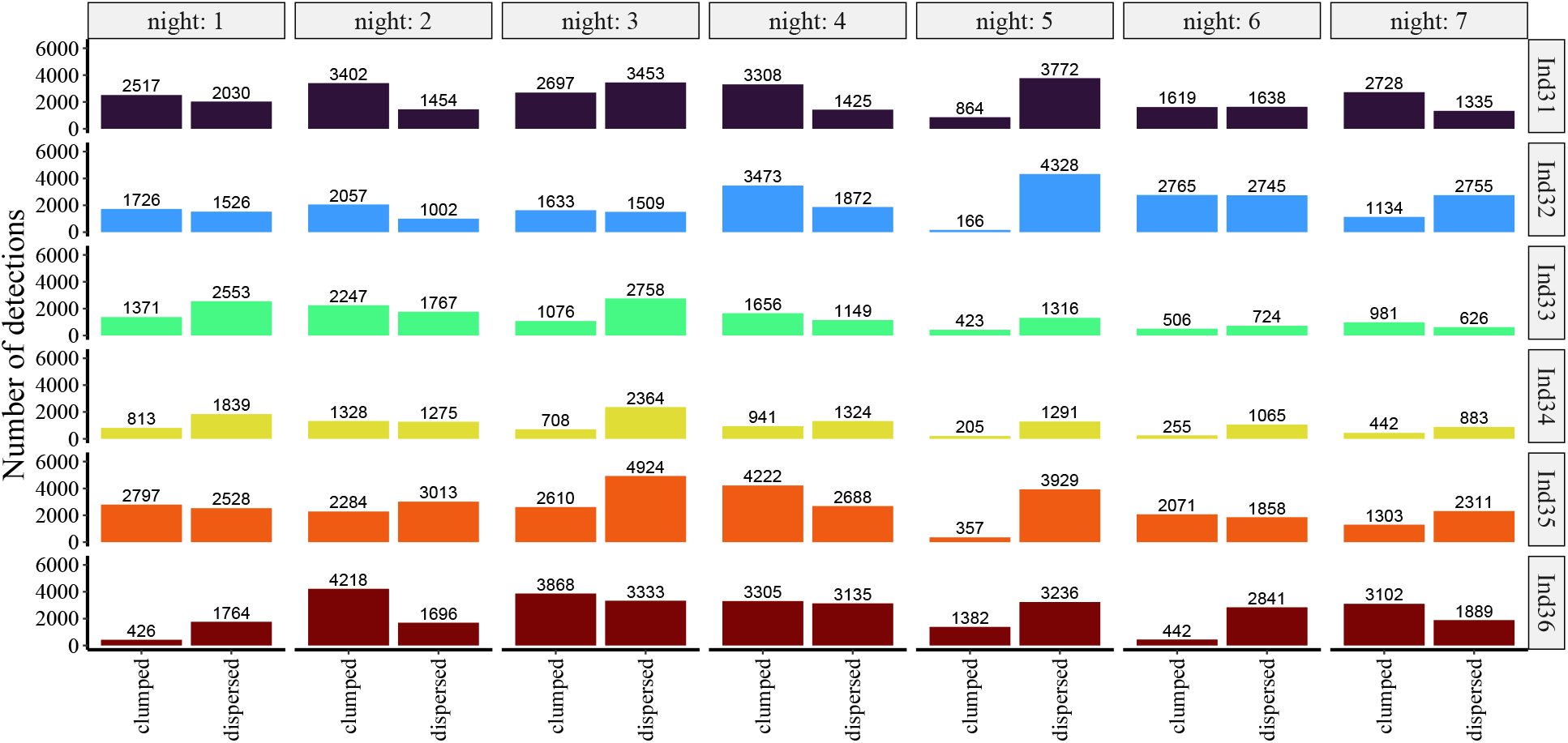
Raw number of detections for all bats in females-only group 1.

**Figure S10:**
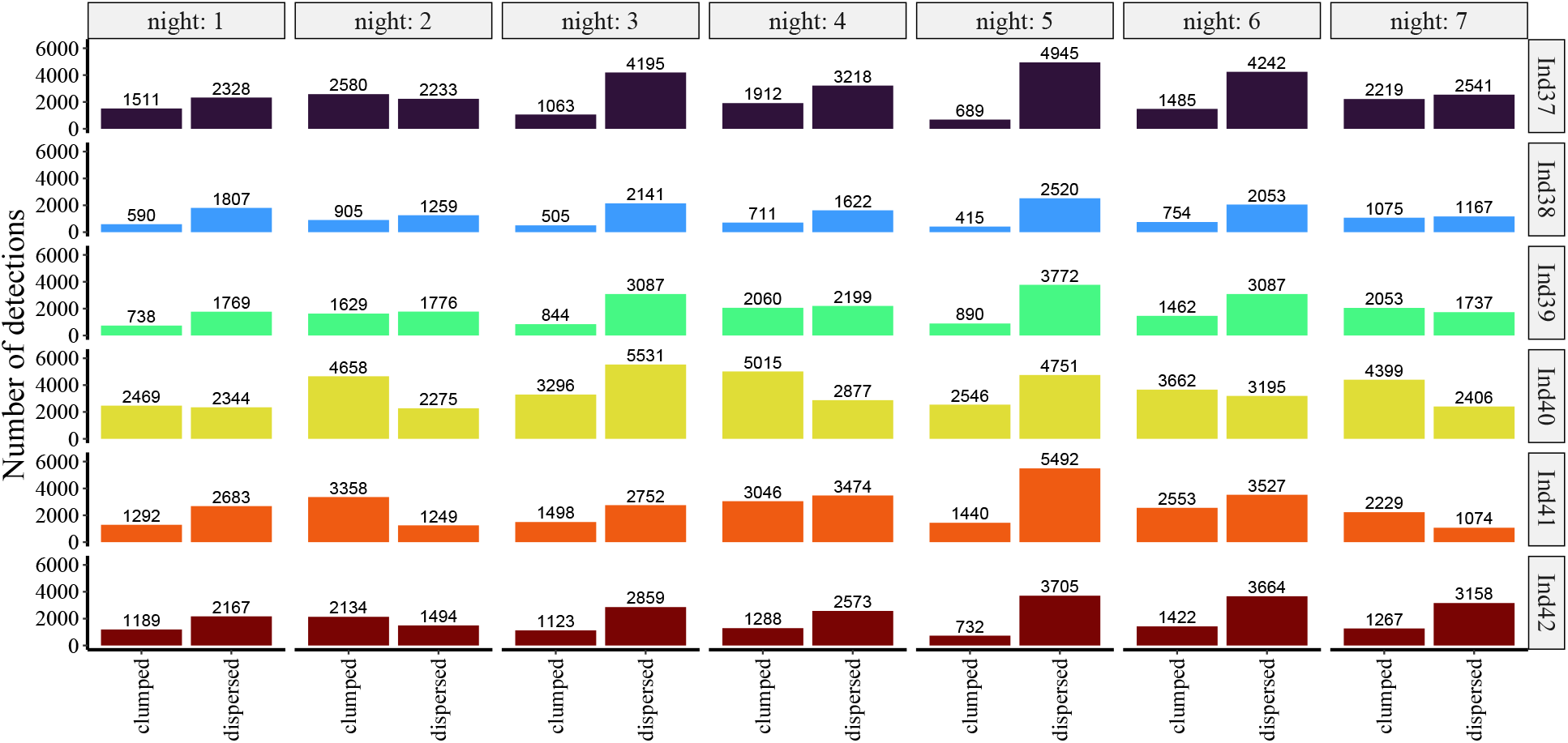
Raw number of detections for all bats in females-only group 2.

**Figure S11:**
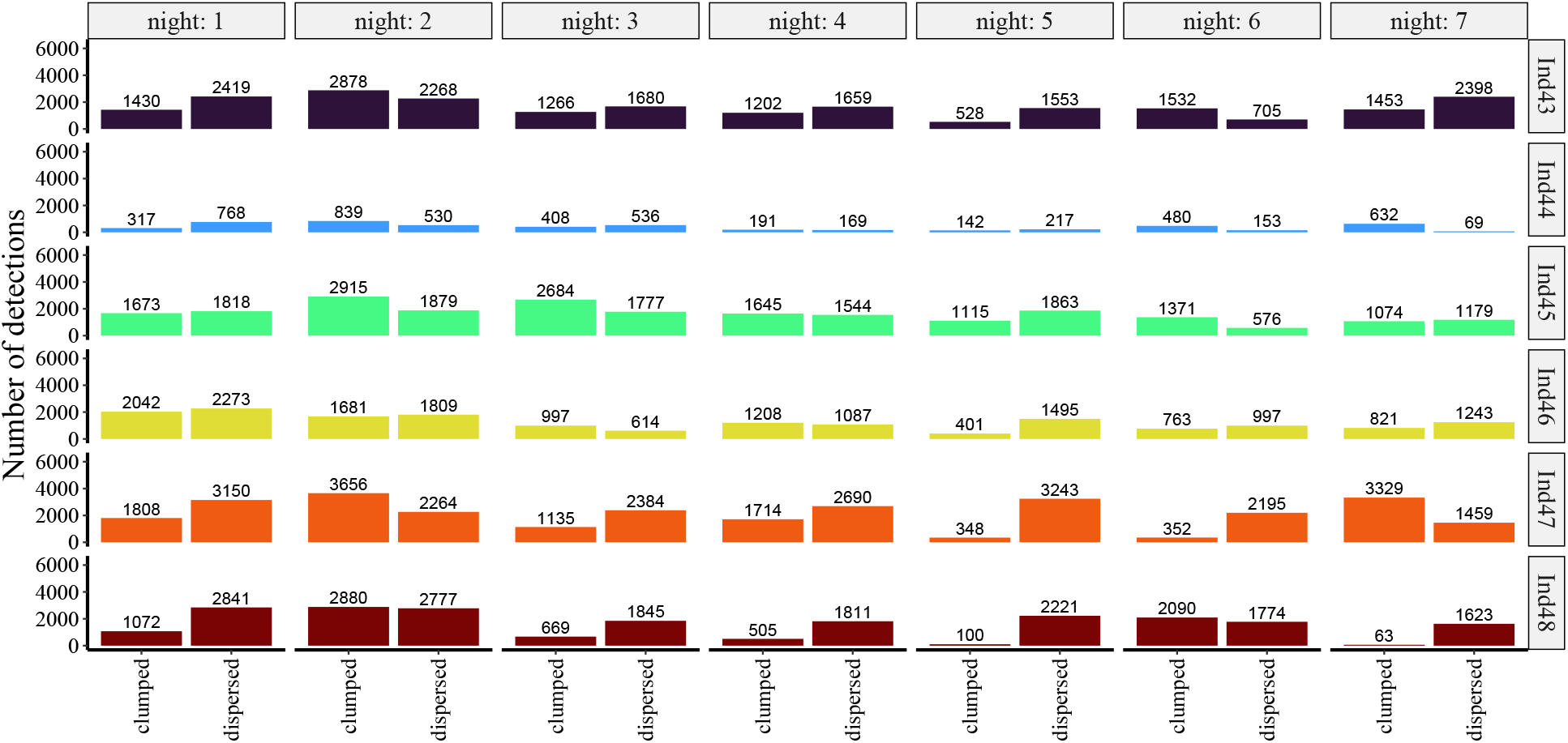
Raw number of detections for all bats in females-only group 3.

**Figure S12:**
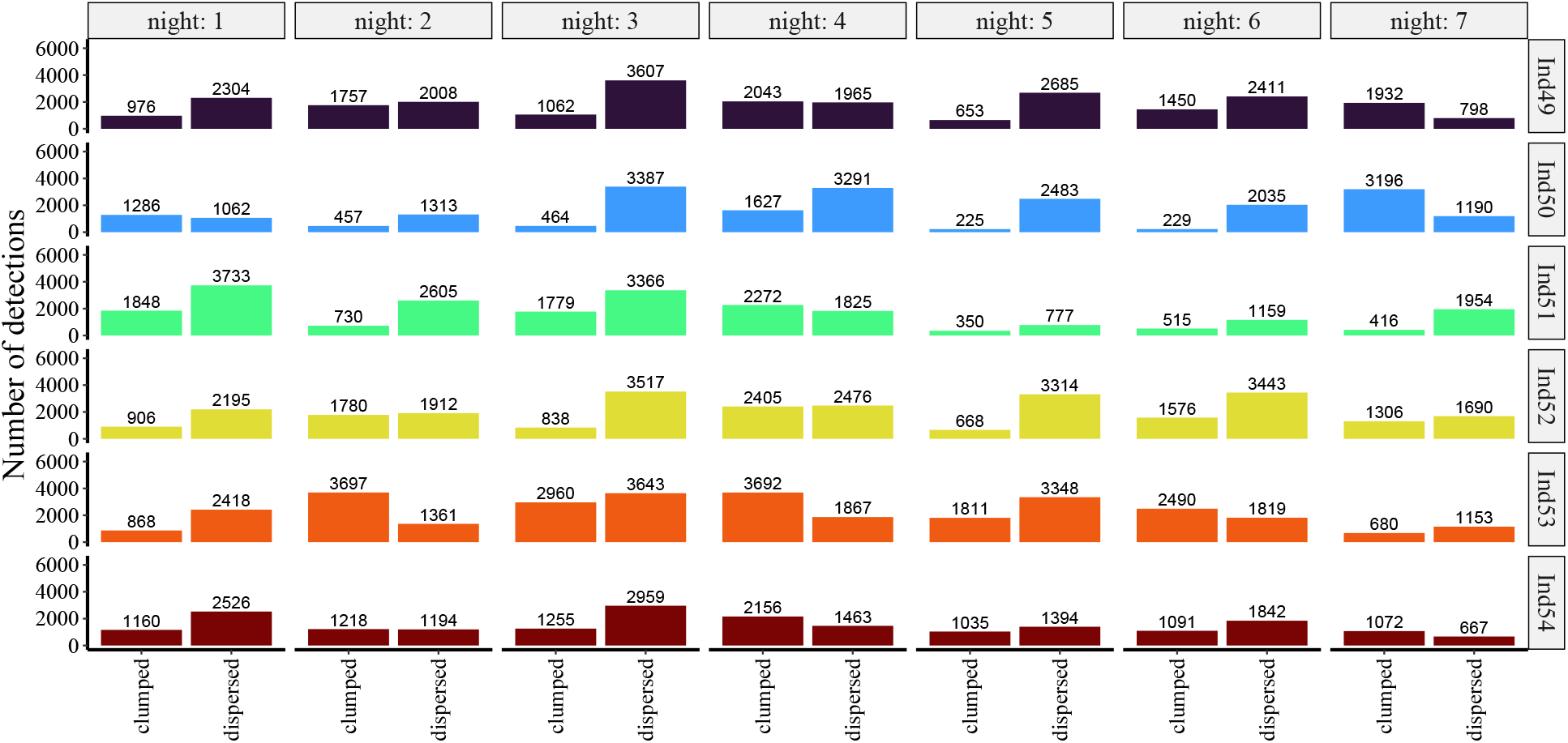
Raw number of detections for all bats in females-only group 4.

**Figure S13:**
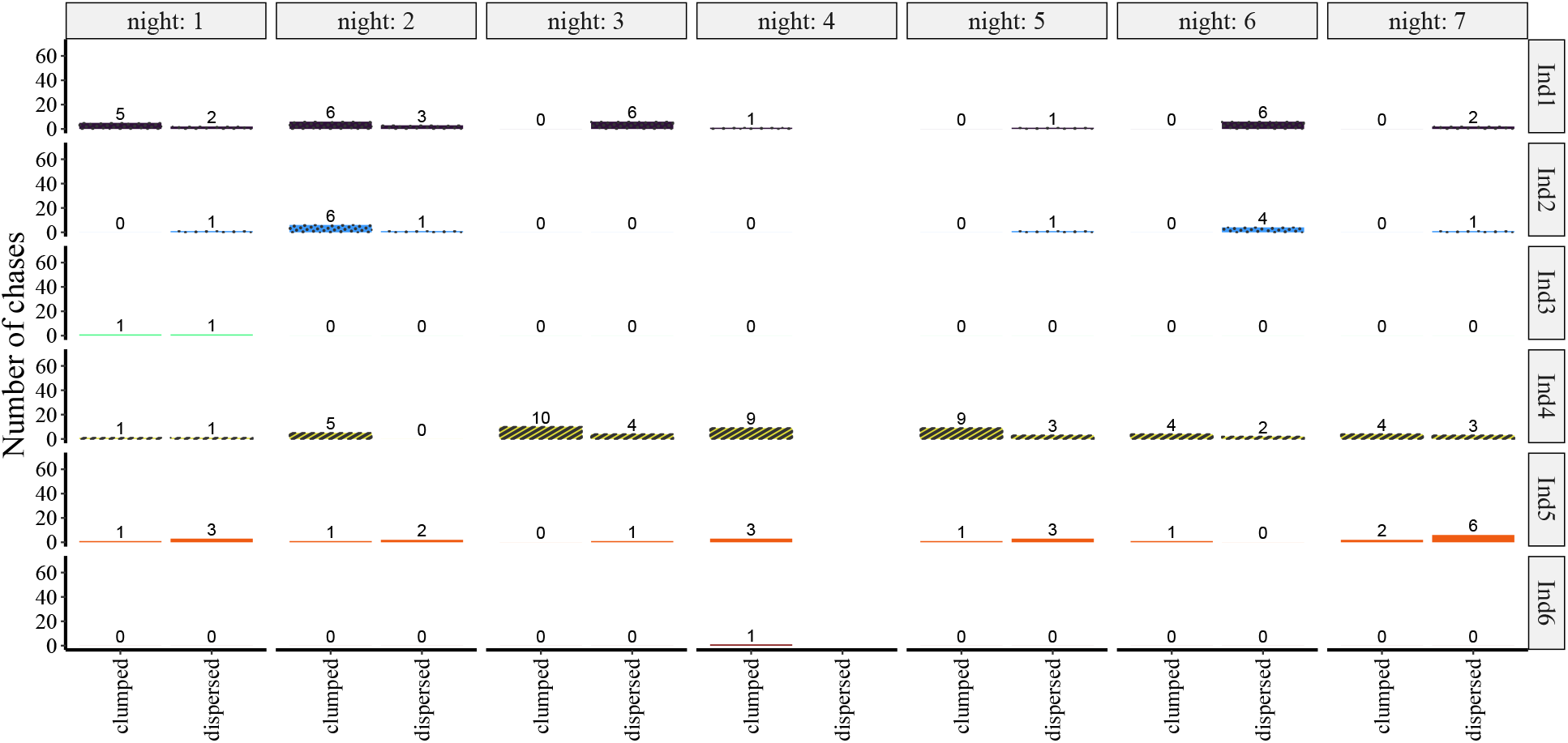
Raw number of chase events for all bats in mixed group 1. Same notation as in Fig. S4, but the colors correspond to different individuals.

**Figure S14:**
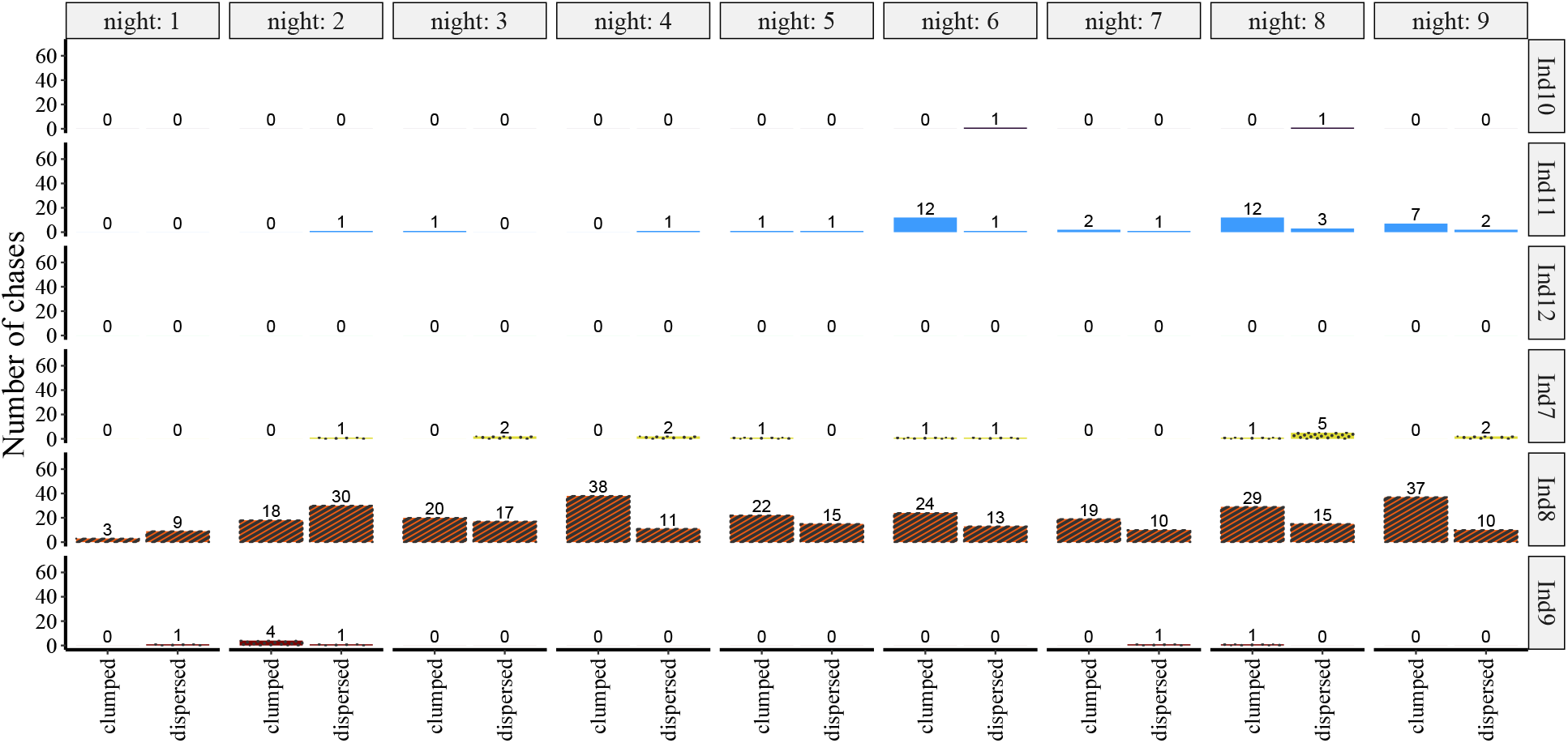
Raw number of chase events for all bats in mixed group 2. Same notation as in Fig. S4, but the colors correspond to different individuals.

**Figure S15:**
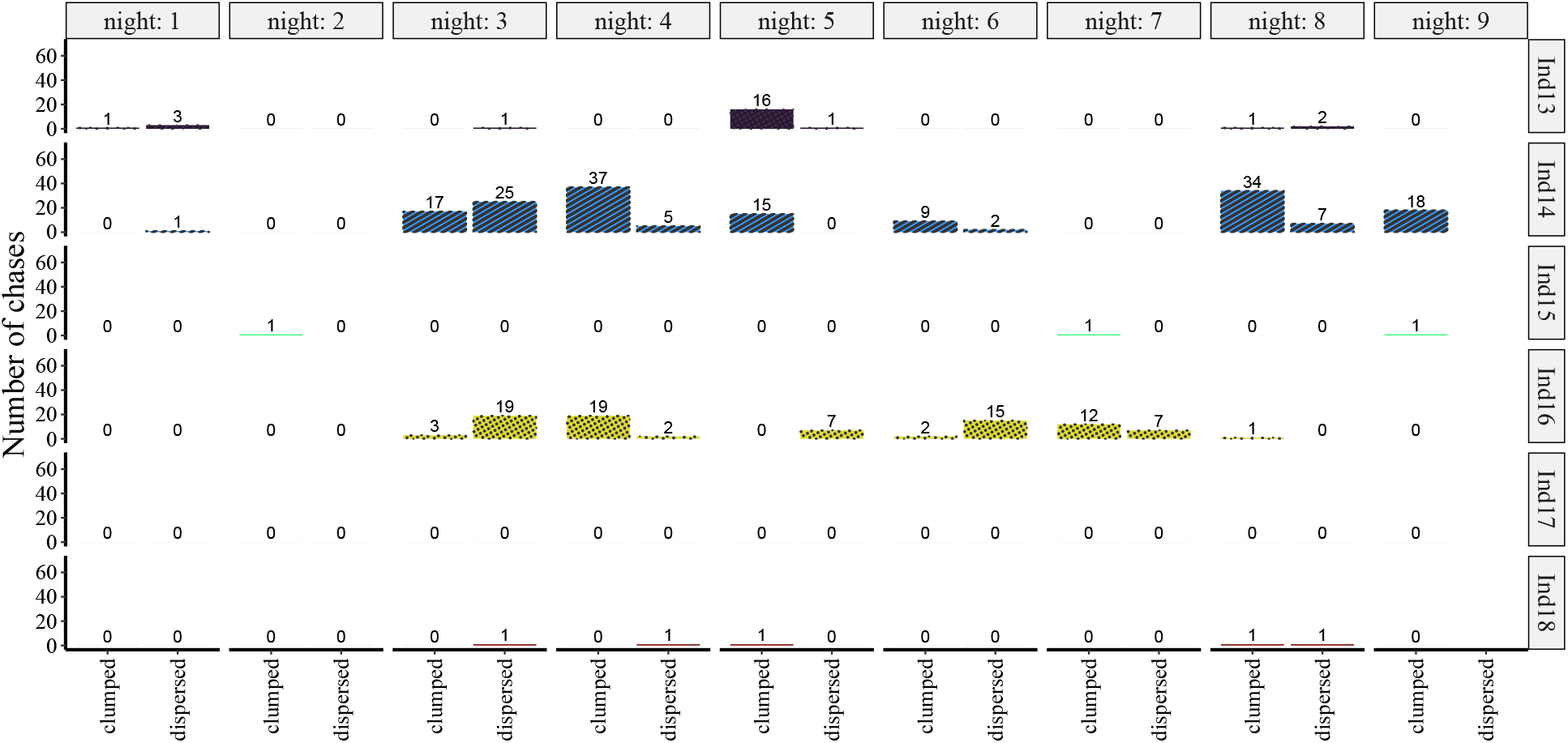
Raw number of chase events for all bats in mixed group 3. Same notation as in Fig. S4, but the colors correspond to different individuals.

**Figure S16:**
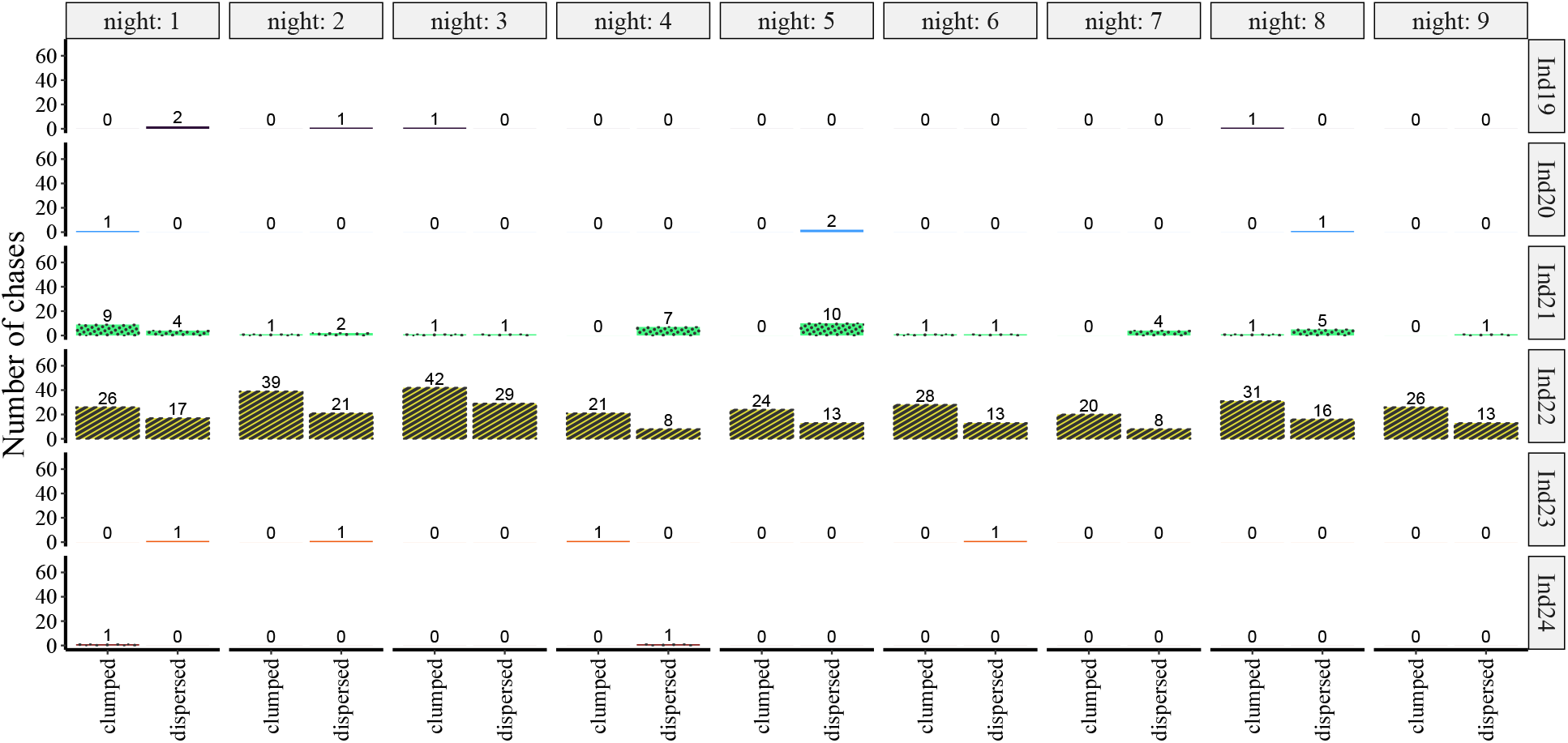
Raw number of chase events for all bats in mixed group 4. Same notation as in Fig. S4, but the colors correspond to different individuals.

**Figure S17:**
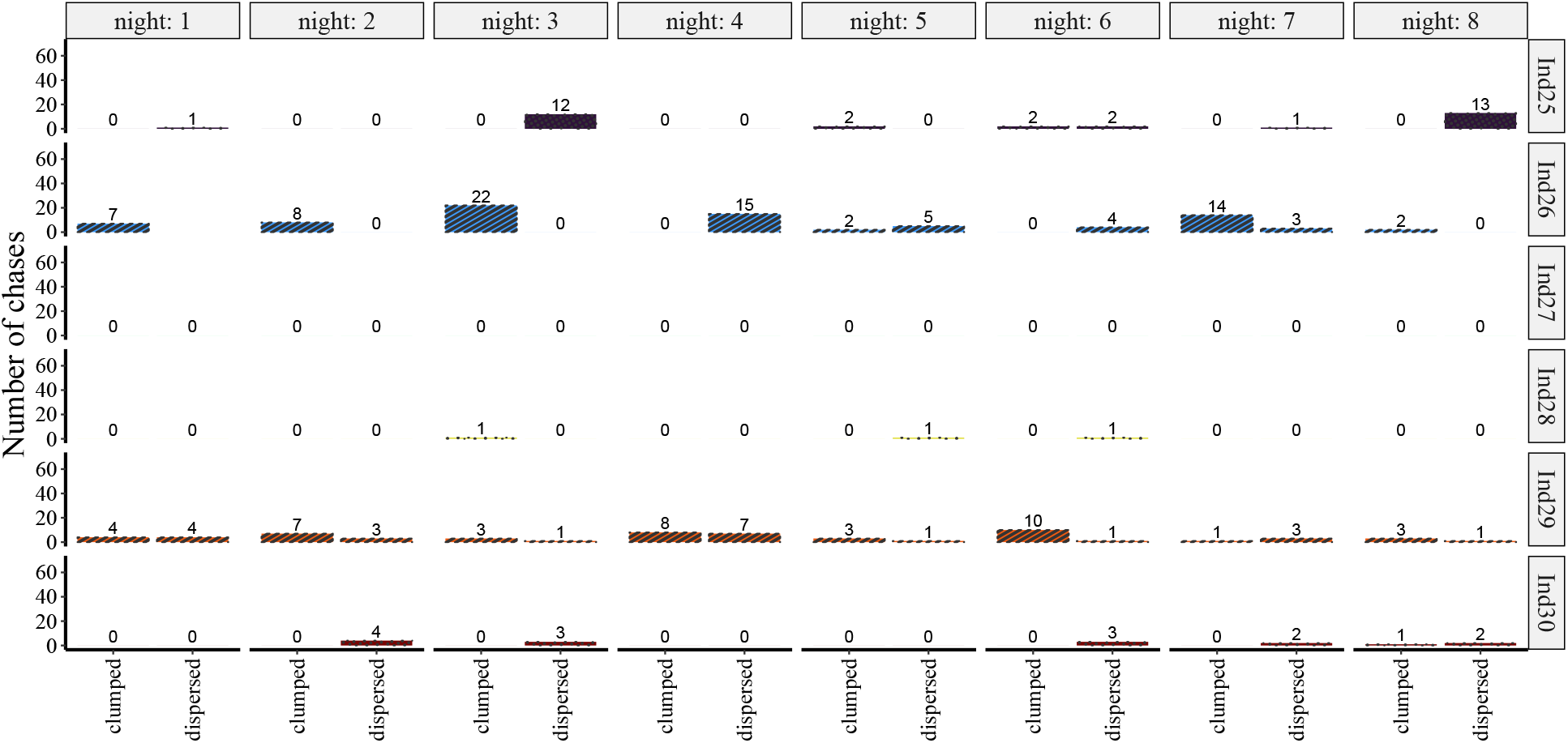
Raw number of chase events for all bats in males-only group. Same notation as in Fig. S4, but the colors correspond to different individuals.

**Figure S18:**
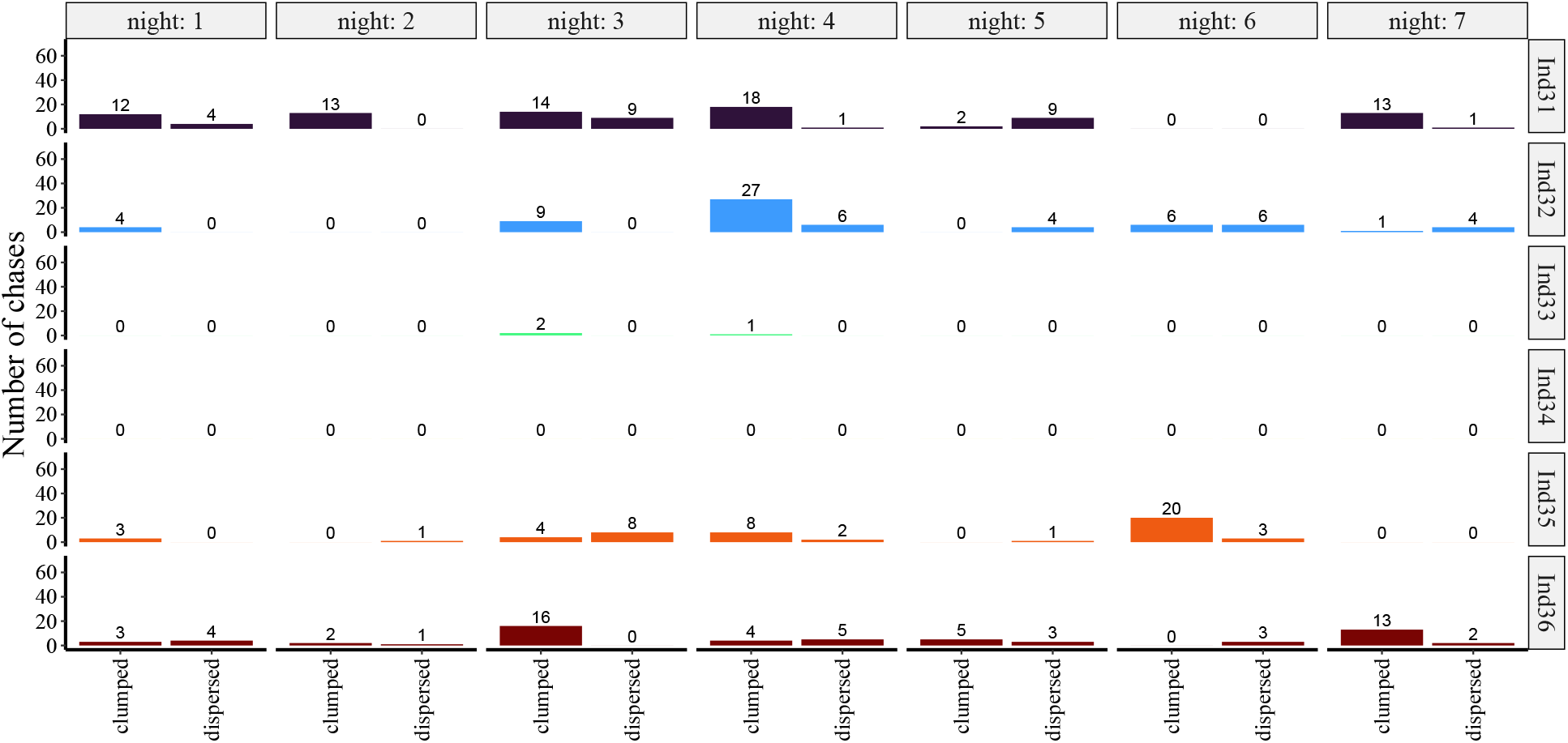
Raw number of chase events for all bats in females-only group 1.

**Figure S19:**
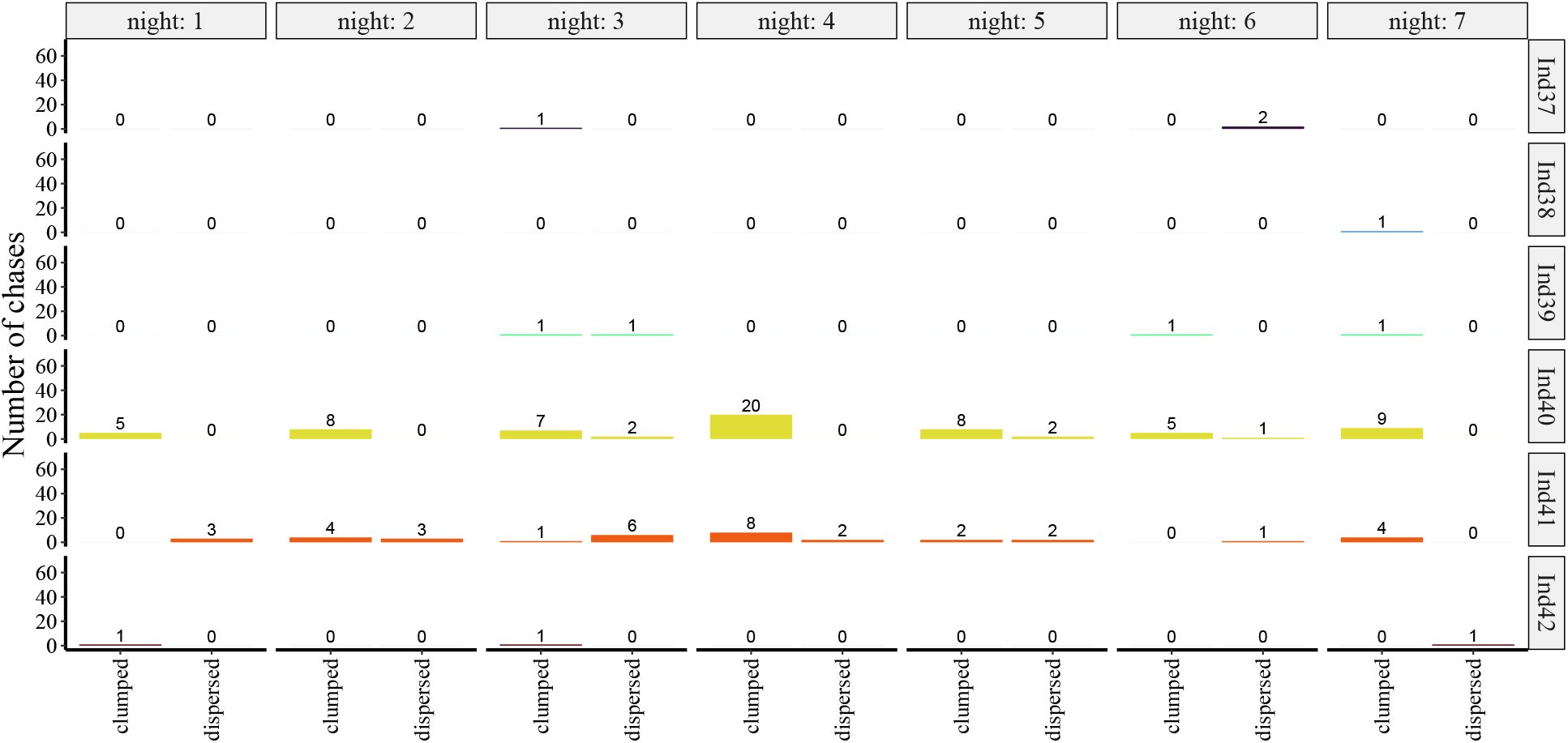
Raw number of chase events for all bats in females-only group 2.

**Figure S20:**
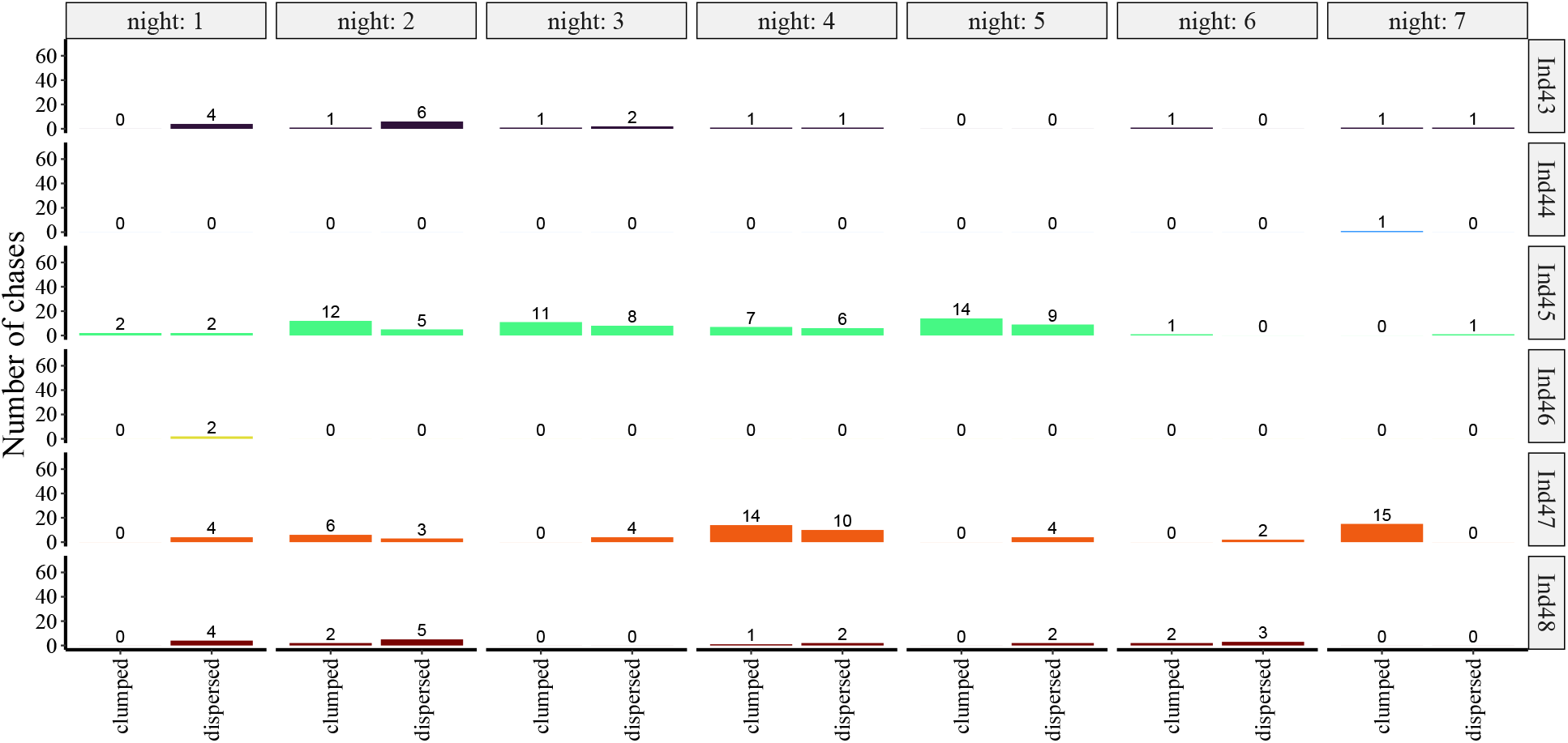
Raw number of chase events for all bats in females-only group 3.

**Figure S21:**
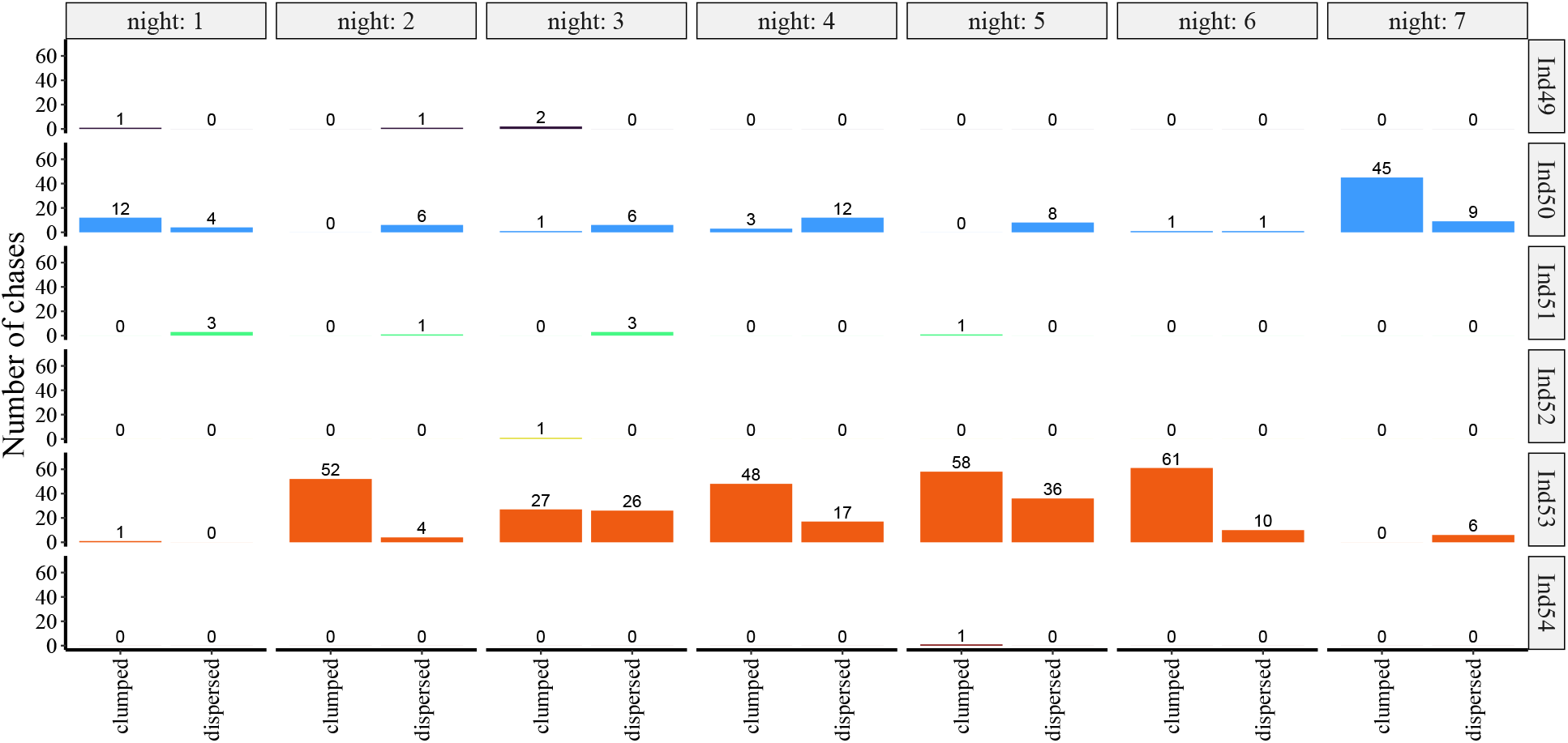
Raw number of chase events for all bats in females-only group 4.

**Figure S22:**
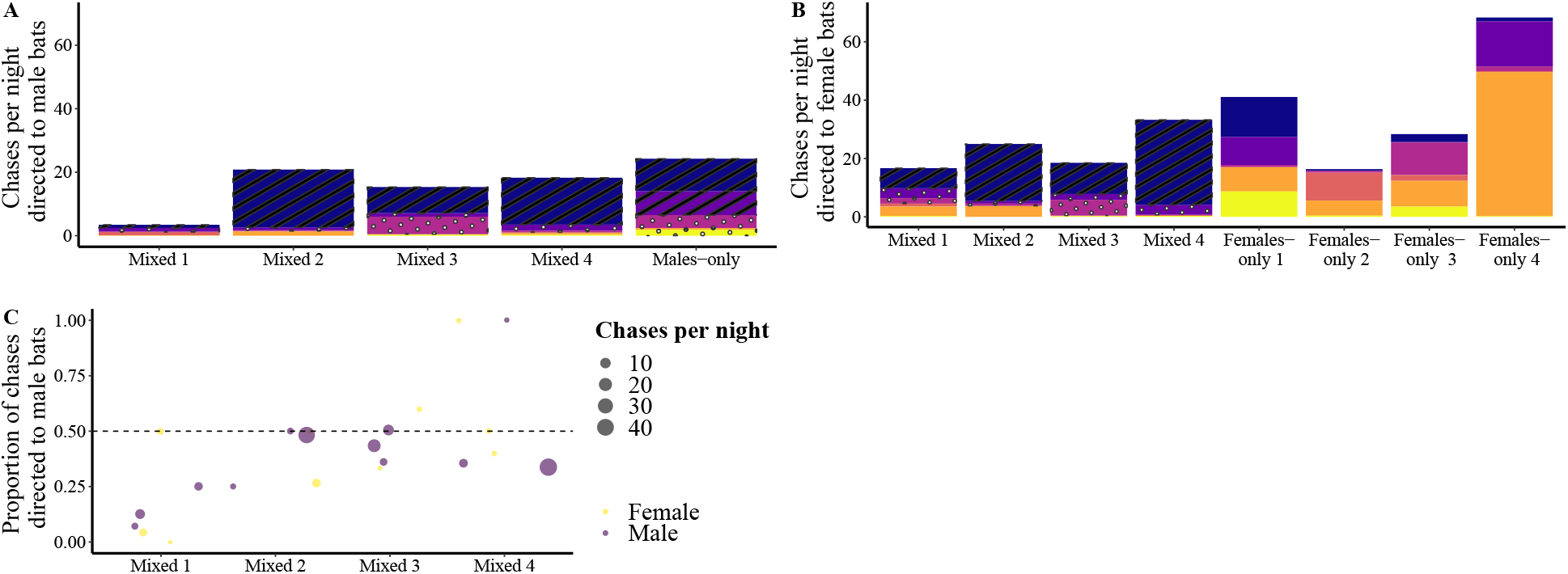
Frequency of chase events directed to bats of either sex. (**A**, **B**) Colored bars give the frequency of chase events per night (total chases divided by number of nights) over the main experiment by each individual from each group (abscissa) towards male (**A**) and female (**B**) bats. The dominant males are shown with stripes and the subordinate males are shown with dots. (**C**) Relative frequency of chase events directed towards male bats for male (purple) and female (yellow) bats from all mixed experimental groups (abscissa). The size of the dots represents the mean frequency of chase events per night.

**Figure S23:**
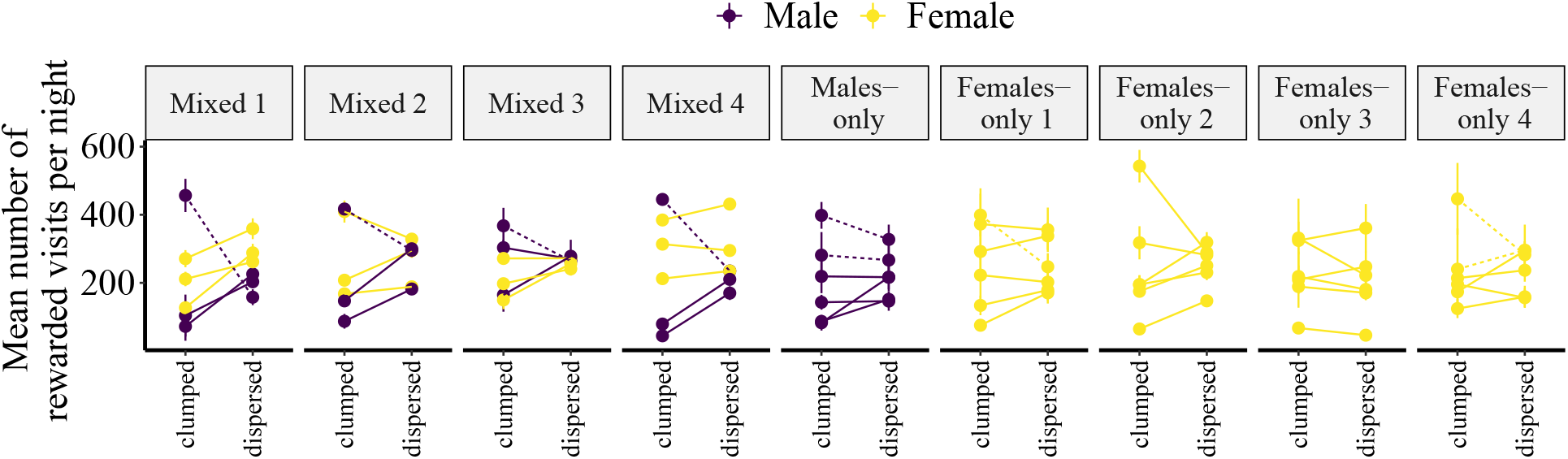
Number of rewarded visits per night (mean ±SE) during the clumped and dispersed reward treatments for male (purple) and female (yellow) bats in each experimental group (panels). Data from the same individuals are connected with lines. Data from individuals classified as “dominant” are connected with dashed lines.

**Figure S24:**
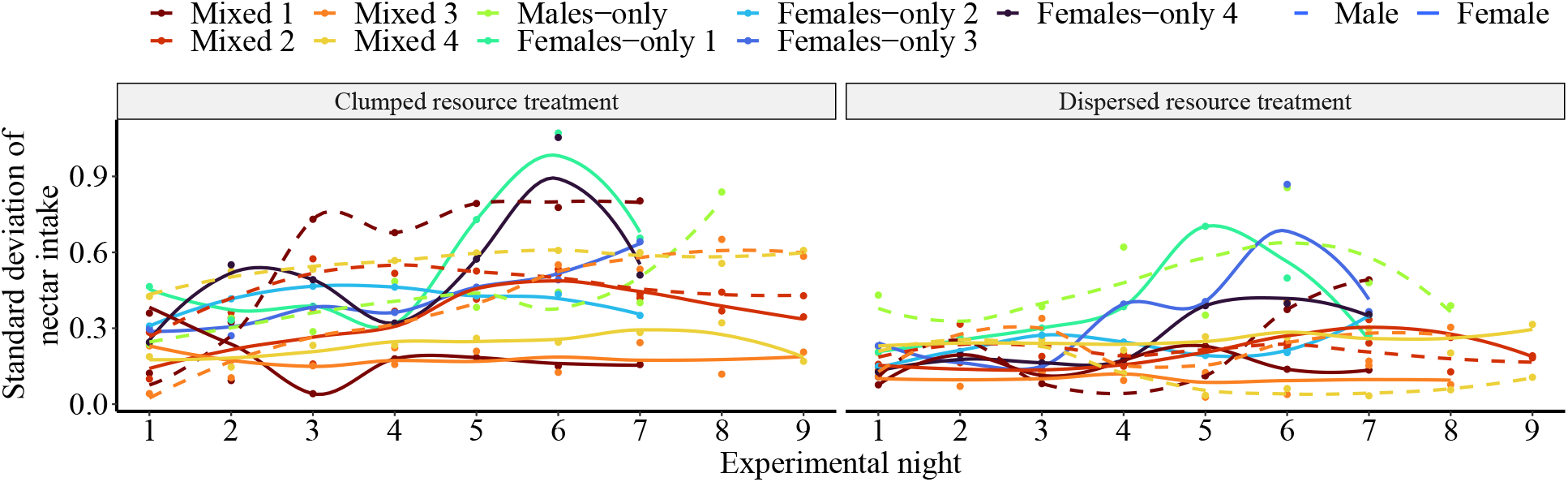
The standard deviation of group nectar consumption was used to measure the between-individual differences in nectar intake. It was calculated for the clumped (left panel) and the dispersed (right panel) resource treatments, separately for males (dashed lines) and females (continuous lines) from each experimental group (different colors). For visualization only, lines give the corresponding fits based on locally weighted scatterplot smoothing (loess). The statistical analysis was based on linear regression (see Methods).

**Figure S25:**
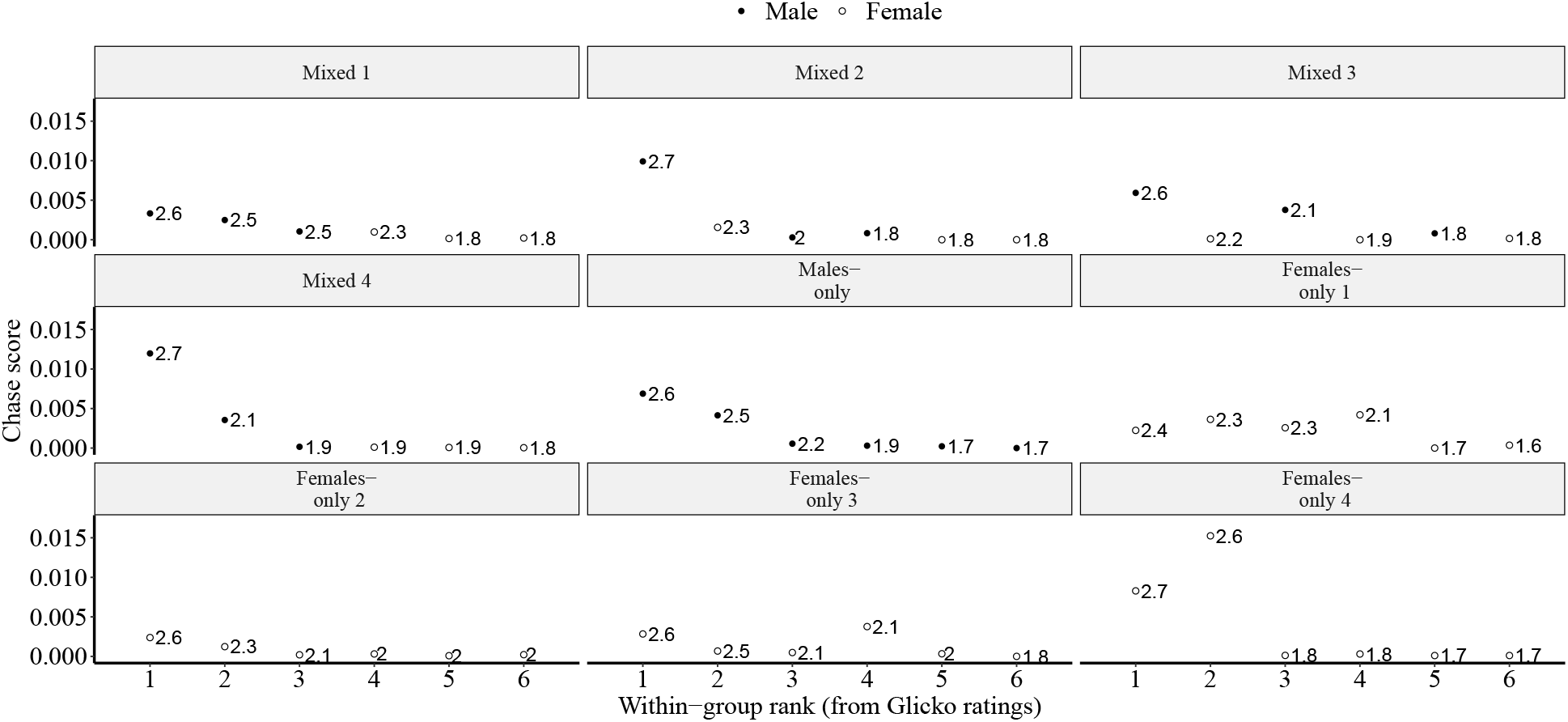
Glicko ratings within the six experimental groups. Over the last two experimental nights, the males (closed symbols) with the highest chase scores were also the individuals with the highest Glicko rating in each group (panels) during the clumped resource treatment. In female-only groups this correspondence was found only in group 2. Numbers at symbols give the Glicko rating in thousands.

**Figure S26:**
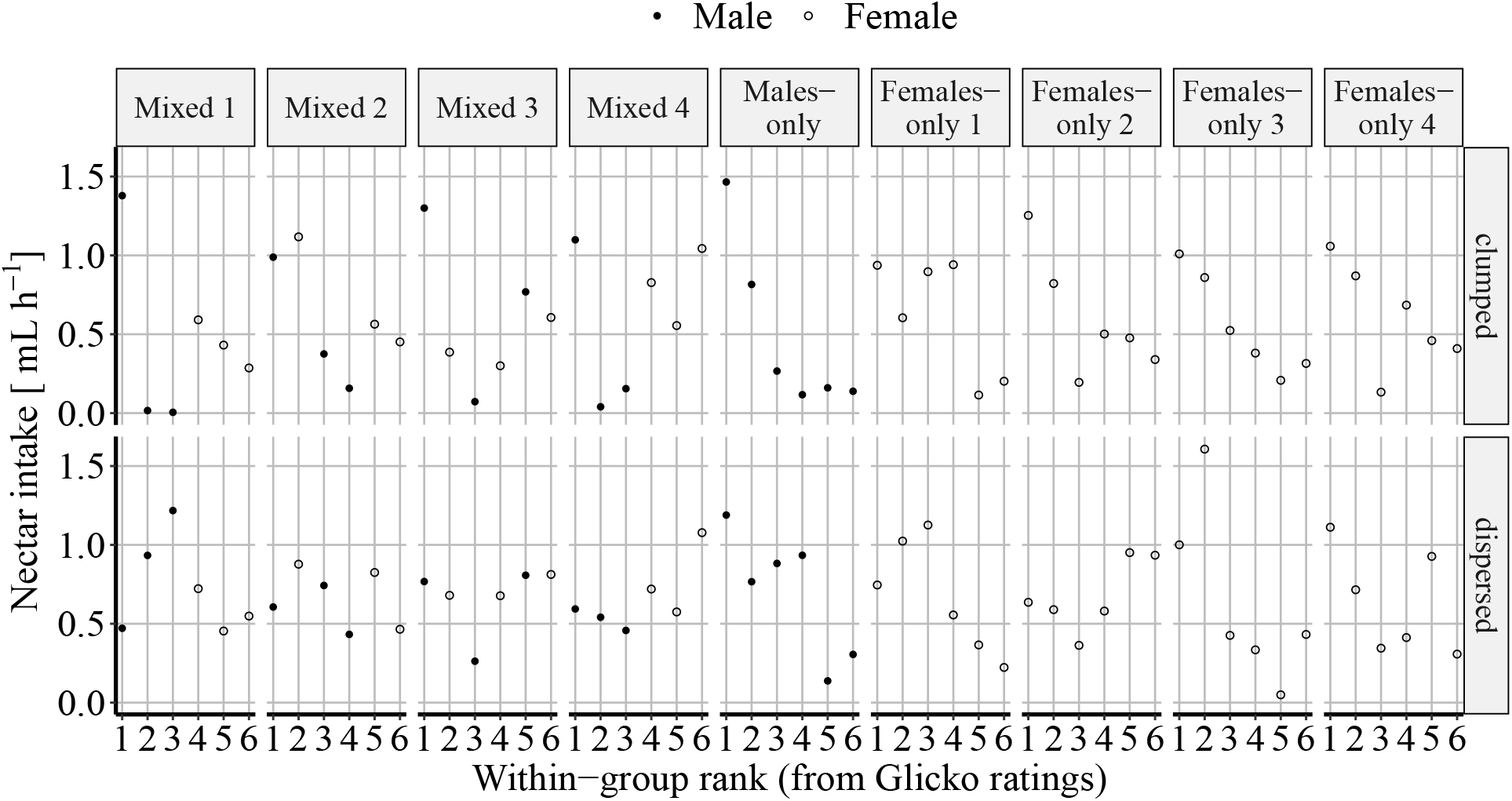
Male bats or female bats in the single-sex groups with the highest Glicko ratings had the highest nectar intake rates during the clumped, but generally not during the dispersed resource treatment.

**Figure S27:**
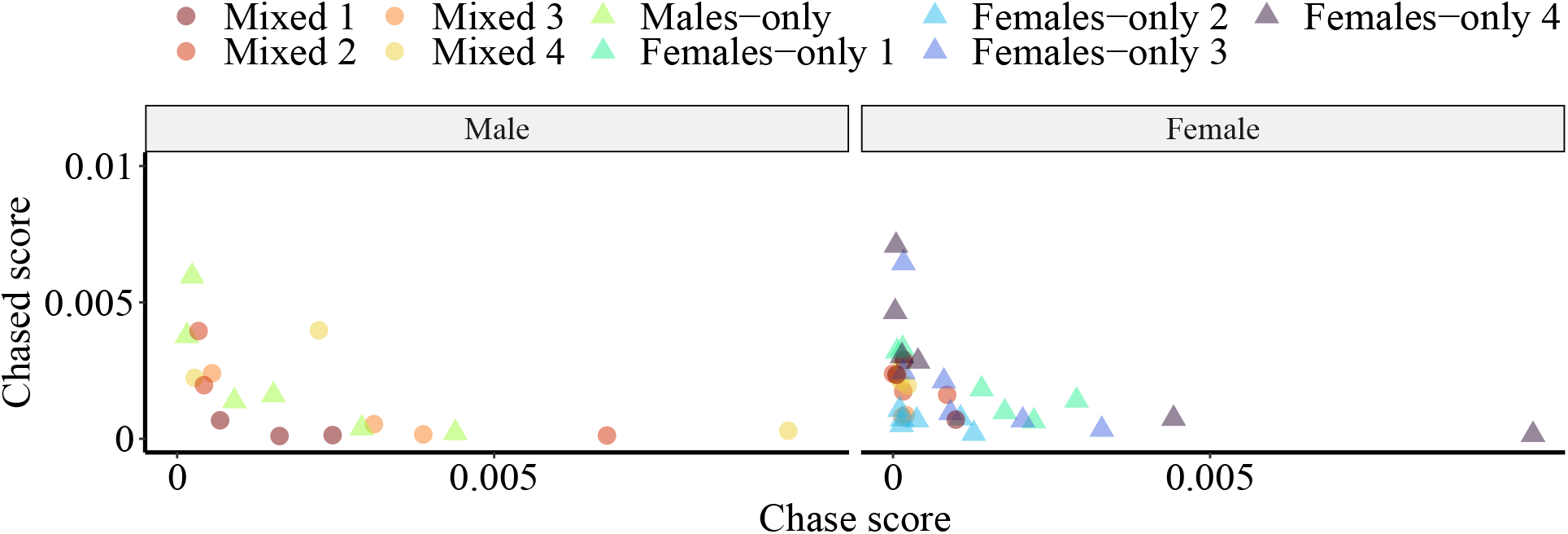
The relationship between chase score and chased score (proportion of chases versus the proportion of being chased out of all detections) for female (right) and male (left) individuals in all experimental groups (different colors). Mixed groups are shown with circles and single-sex groups, with triangles.

**Table S1:**
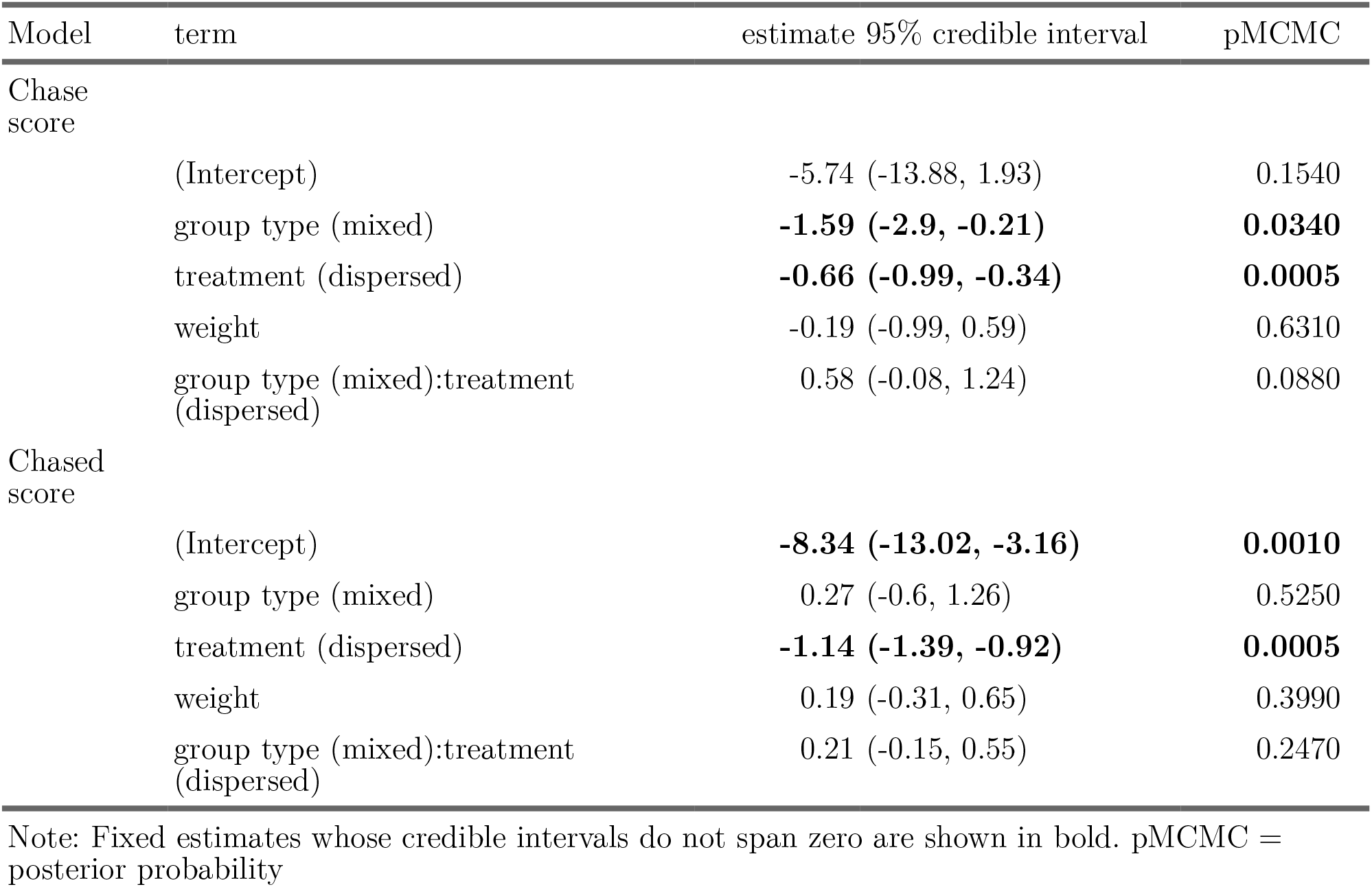
Summary of fixed effects from generalized linear mixed-effects models of chasing frequency and the frequency of being chased for female bats only.

| Model | term | estimate 95% credible interval | pMCMC |
| --- | --- | --- | --- |
| Chase score | (Intercept) | -5.74 (-13.88, 1.93) | 0.1540 |
|  | group type (mixed) | **-1.59 (-2.9, -0.21)** | **0.0340** |
|  | treatment (dispersed) | **-0.66 (-0.99, -0.34)** | **0.0005** |
|  | weight | -0.19 (-0.99, 0.59) | 0.6310 |
|  | group type (mixed):treatment (dispersed) | 0.58 (-0.08, 1.24) | 0.0880 |
| Chased score | (Intercept) | **-8.34 (-13.02, -3.16)** | **0.0010** |
|  | group type (mixed) | 0.27 (-0.6, 1.26) | 0.5250 |
|  | treatment (dispersed) | **-1.14 (-1.39, -0.92)** | **0.0005** |
|  | weight | 0.19 (-0.31, 0.65) | 0.3990 |
|  | group type (mixed):treatment (dispersed) | 0.21 (-0.15, 0.55) | 0.2470 |

Note: Fixed estimates whose credible intervals do not span zero are shown in bold. pMCMC = posterior probability

**Table S2:** Summary of fixed effects from a generalized linear mixed-effects model of the standard deviation of nectar intake over time for female bats only.

| term | estimate 95% credible interval | pMCMC |
| --- | --- | --- |
| (Intercept) | **0.50 (0.41, 0.6)** | **0.000** |
| group type (mixed) | **-0.27 (-0.4, -0.14)** | **0.003** |
| treatment (dispersed) | **-0.18 (-0.24, -0.11)** | **0.000** |
| night | **0.05 (0.01, 0.1)** | **0.030** |
| group type (mixed):treatment (dispersed) | **0.12 (0.04, 0.2)** | **0.007** |
| group type (mixed):night | -0.05 (-0.11, 0.01) | 0.116 |
| treatment (dispersed):night | -0.01 (-0.04, 0.02) | 0.573 |
| group type (mixed):treatment (dispersed):night | 0.01 (-0.03, 0.05) | 0.648 |

Note: Fixed estimates whose credible intervals do not span zero are shown in bold. pMCMC = posterior probability

**Figure S28:**
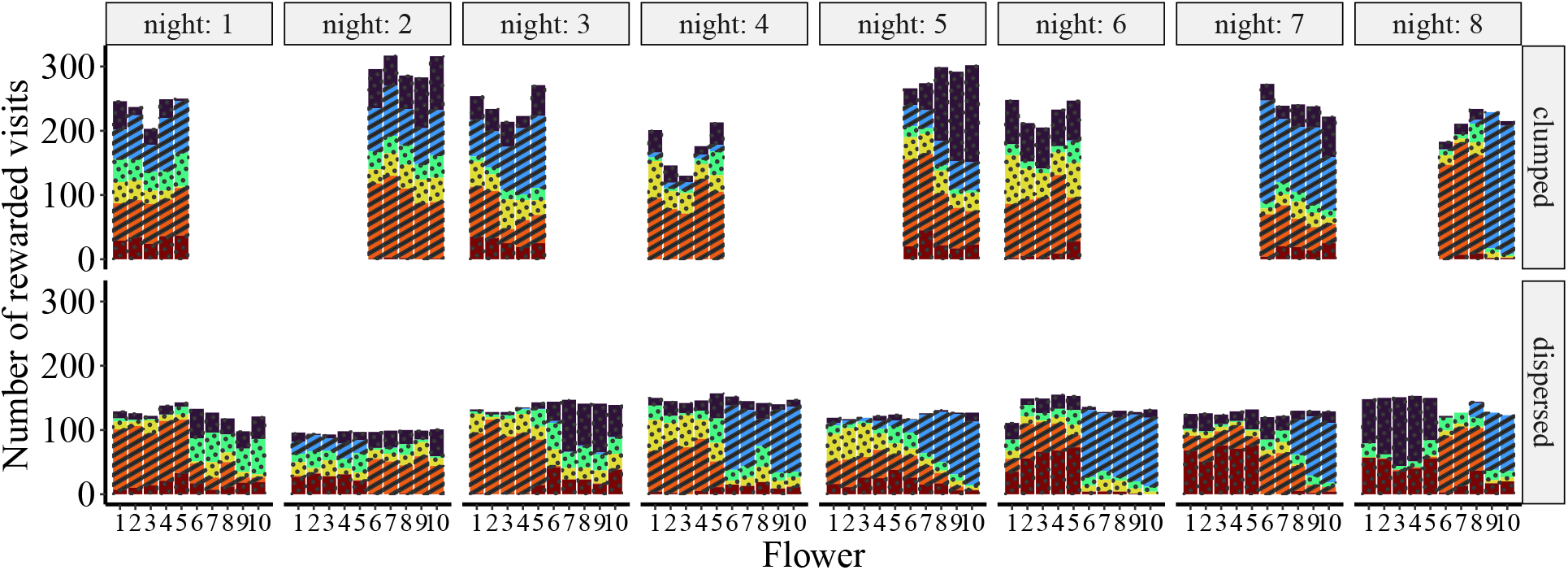
Distribution of rewarded visits across flowers for the six bats in the males-only group. The colored bars give the number of rewarded visits of each individual at the ten flowers during the clumped (top) and dispersed (bottom) resource treatments for each experimental night (columns). The dominant males are shown with stripes and the subordinate males are shown with dots. This was the only group with two males behaving as dominant. On the last night, rather than sharing all flowers within the defended patch, the dominant males partitioned the patch into two subpatches, with each bat defending its own partition.

**Figure S29:**
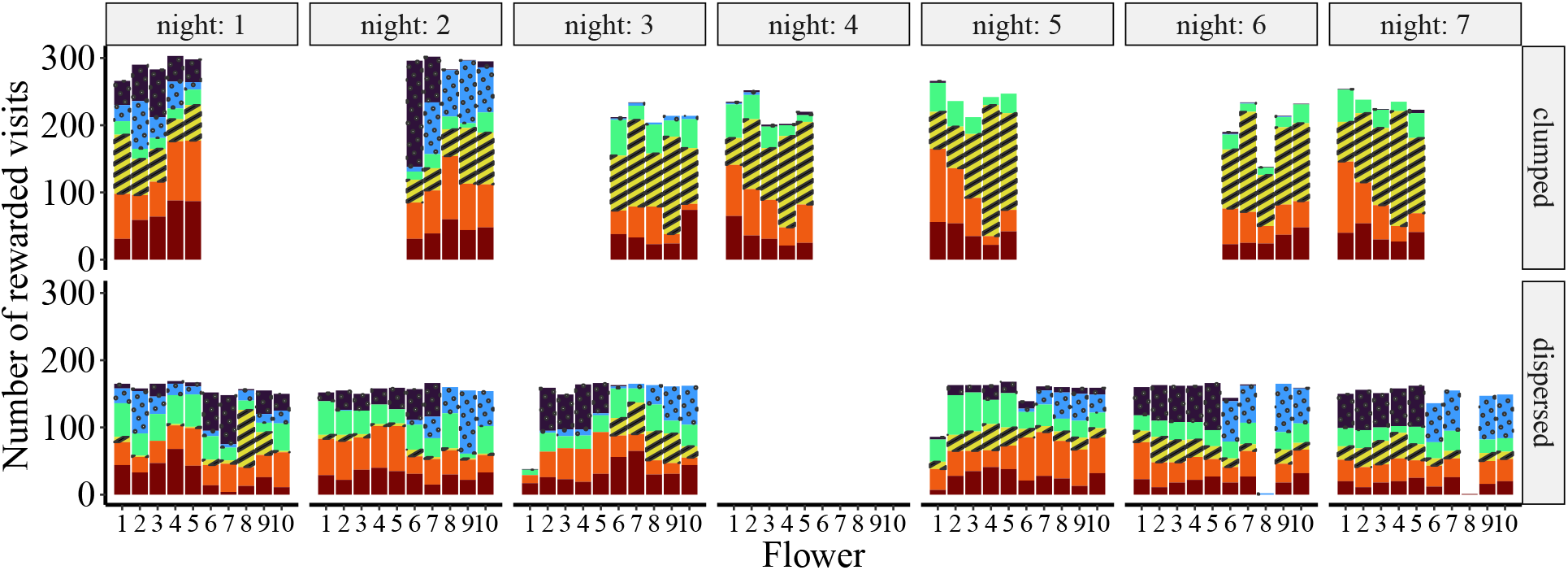
Distribution of rewarded visits across flowers for the six bats in mixed group 1. The colored bars give the number of rewarded visits of each individual at the ten flowers during the clumped (top) and dispersed (bottom) resource treatments for each experimental night (columns). The dominant male is shown with stripes, the subordinate males are shown with dots, and the females are shown with solid bars. Due to a technical malfunction on night 4, there were no rewards delivered in the dispersed resource treatment and the data were excluded from analysis.

**Figure S30:**
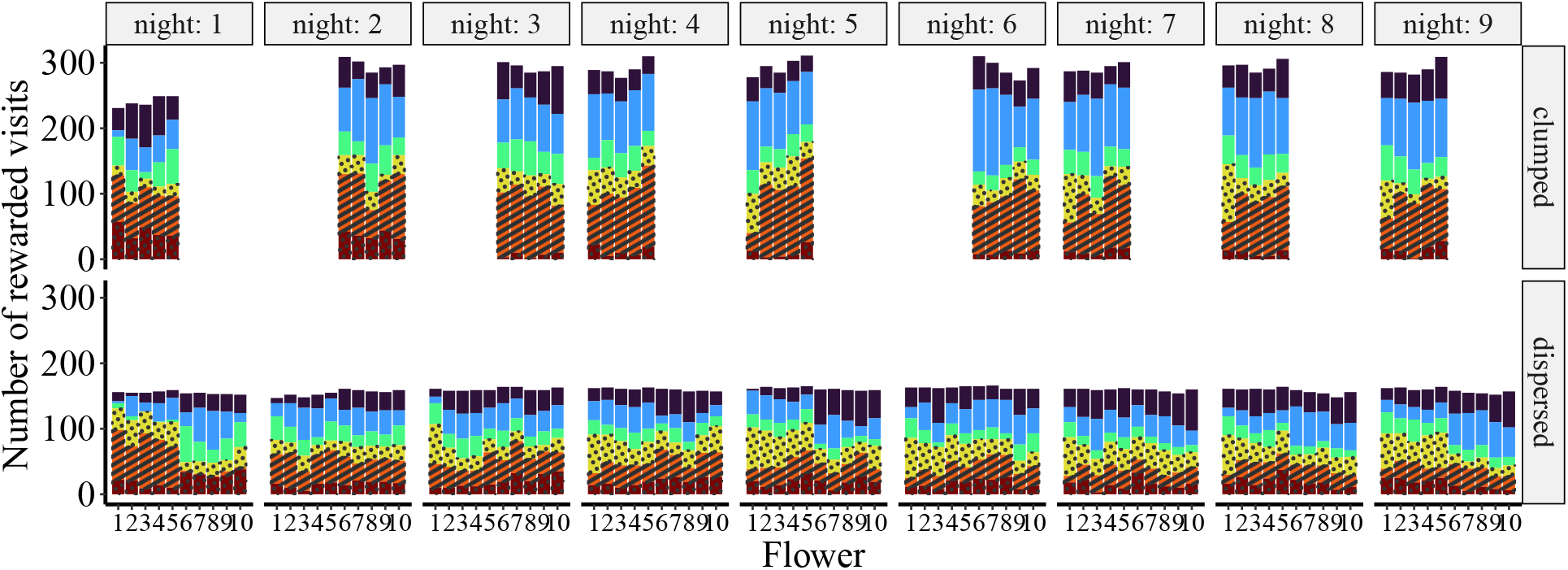
Distribution of rewarded visits across flowers for the six bats in mixed group 2. Same notation as in Fig. S29, but the colors correspond to different individuals.

**Figure S31:**
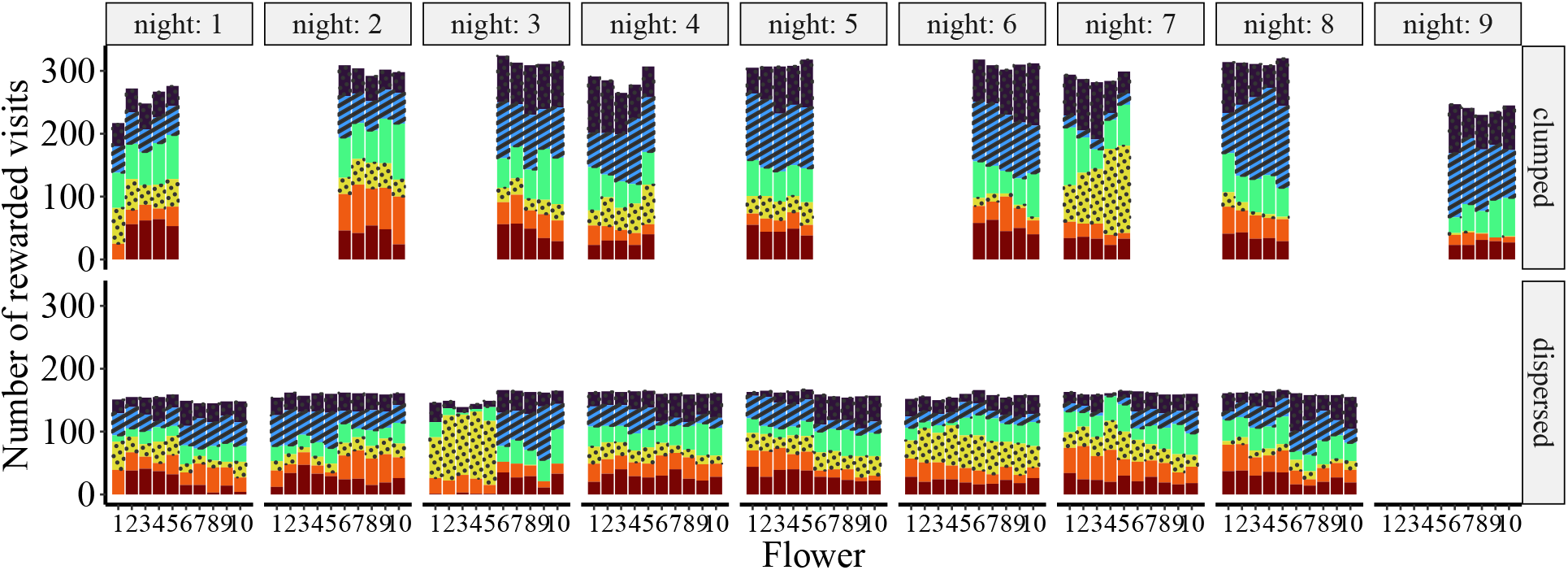
Distribution of rewarded visits across flowers for the six bats in mixed group 3. Same notation as in Fig. S29, but the colors correspond to different individuals. Due to a technical malfunction on night 9, there were no rewards delivered in the dispersed resource treatment and the data were excluded from analysis.

**Figure S32:**
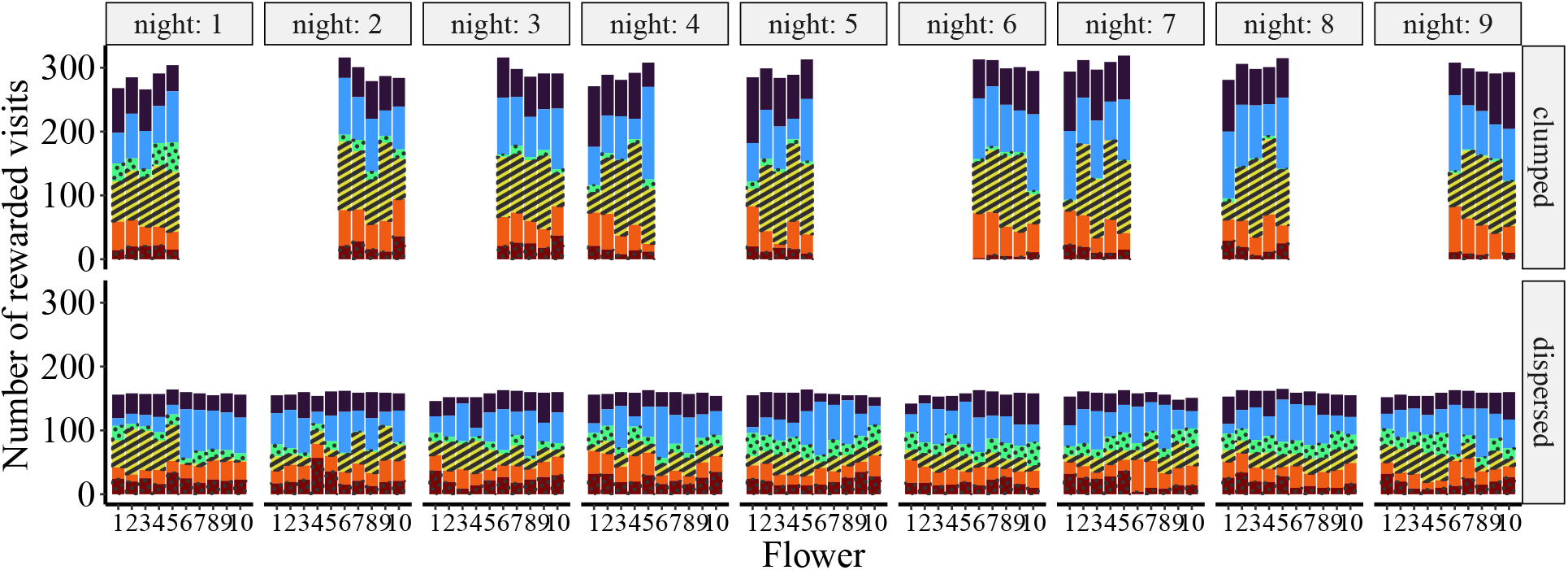
Distribution of rewarded visits across flowers for the six bats in mixed group 4. Same notation as in Fig. S29, but the colors correspond to different individuals.

**Figure S33:**
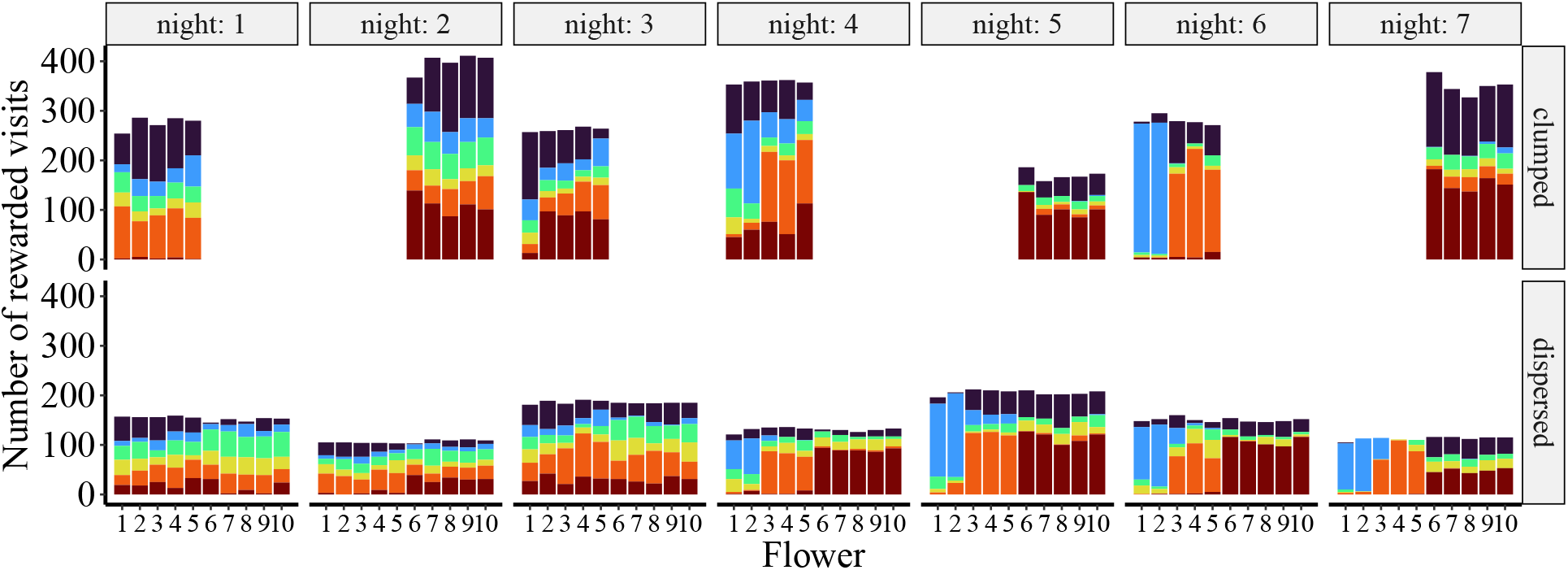
Distribution of rewarded visits across flowers for the six bats in the females-only group 1. Same notation as in Fig. S29, but the colors correspond to different individuals.

**Figure S34:**
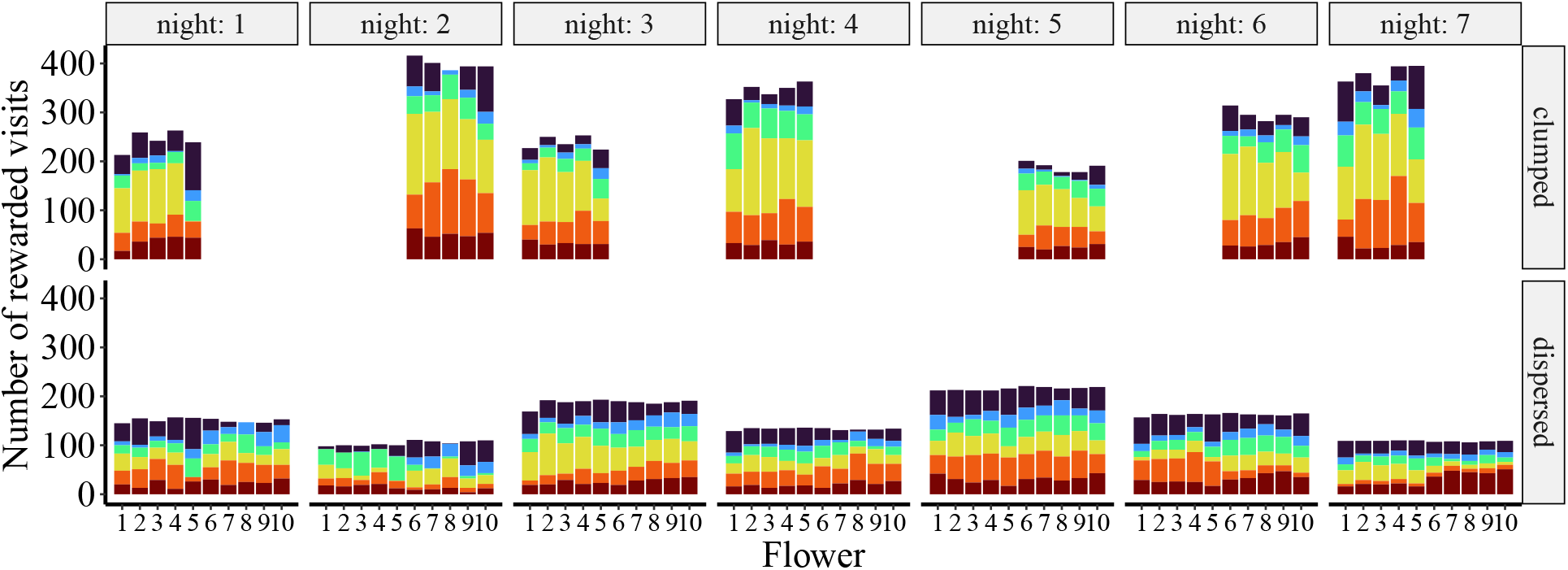
Distribution of rewarded visits across flowers for the six bats in the females-only group 2. Same notation as in Fig. S29, but the colors correspond to different individuals.

**Figure S35:**
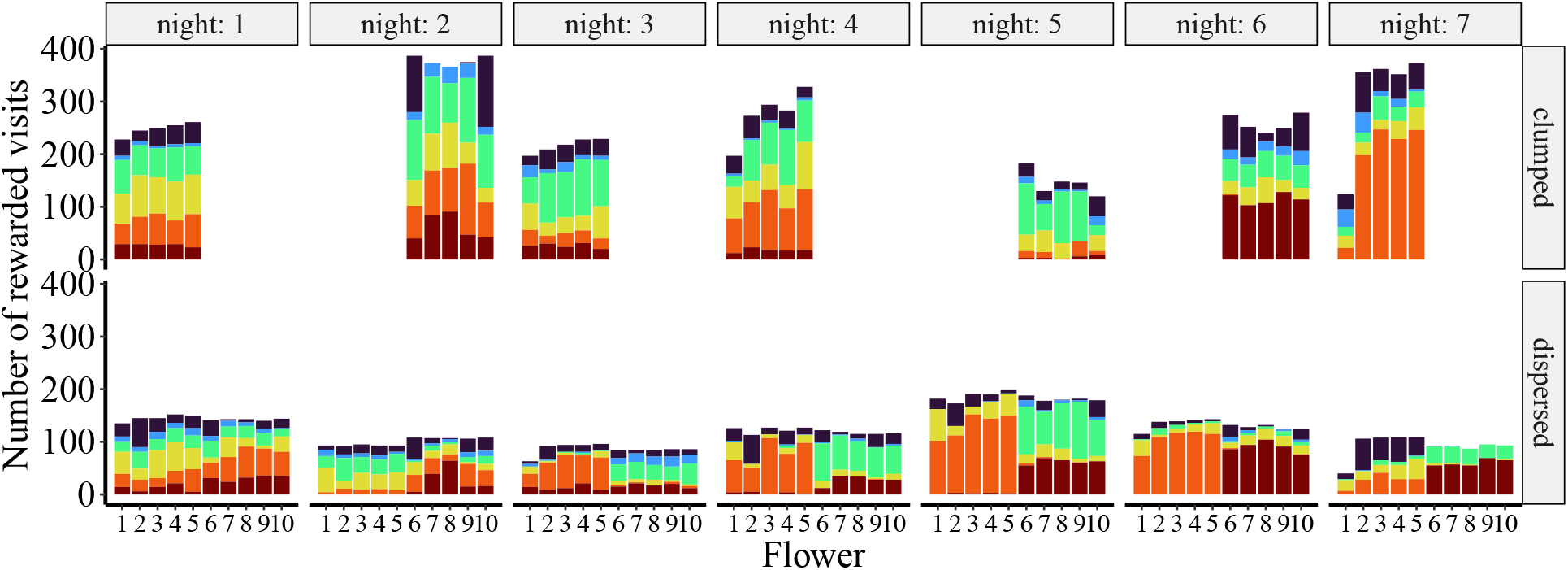
Distribution of rewarded visits across flowers for the six bats in the females-only group 3. Same notation as in Fig. S29, but the colors correspond to different individuals.

**Figure S36:**
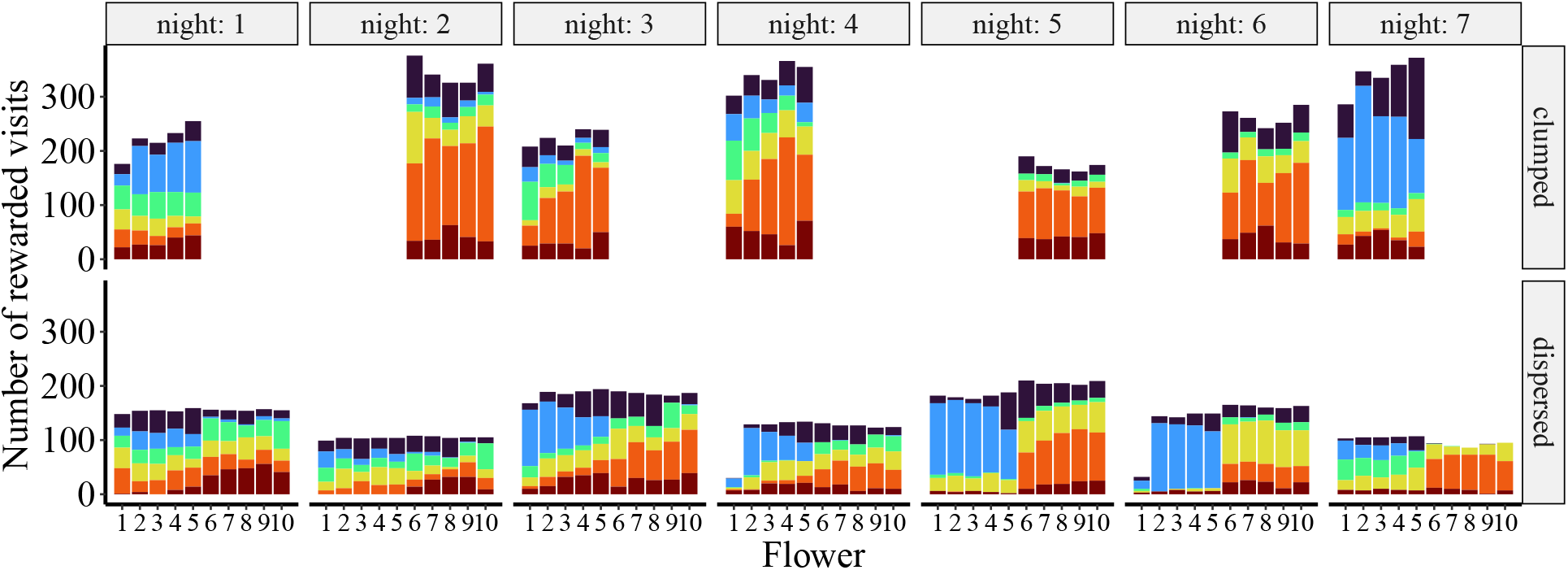
Distribution of rewarded visits across flowers for the six bats in the females-only group 4. Same notation as in Fig. S29, but the colors correspond to different individuals.

## Authors’ contributions

S.W. Conceptualization, Methodology, Software, Data collection, Formal Analysis, Video Analysis, Writing— original draft. V.N. Conceptualization, Methodology, Software, Formal Analysis, Data curation, Writing— review and editing, Visualization, Supervision, Project Administration.

Y.W. Conceptualization, Resources, Methodology, Software (data acquisition), Writing—review and editing, Supervision, Funding.

## Acknowledgements

We thank Katja Frei and Peggy Hoffmann for assistance with the main experiments and animal keeping, Alexej Schatz for programming of the control software, and Peter Spende, Francesco Bagorda, and Waldemar Krzok for help with the experimental hardware. We also thank Marion Rivalan and Lucille Alonso for a fruitful discussion on the analysis of social behavior. We also thank Theo Bakker and three anonymous reviewers for their helpful comments and suggestions for improving the manuscript.

## Data Availability

All data and code are available in the Zenodo repository: https://doi.org/10.5281/zenodo.7581235.

